# Structural and biophysical insights into the unique RNase YicC

**DOI:** 10.64898/2026.09.15.751841

**Authors:** Ruoxi Wu, Harsha Venkatesh, Carisse Lansiquot, Sarah A. Barnes, Afrooz Golestanian, Olga Rechkoblit, Alisha N. Jones, David H. Bechhofer, Michael B. Lazarus

## Abstract

RNA cleavage and processing are highly conserved regulatory mechanisms across bacteria. We recently discovered a novel family of RNases called YicC, conserved across all bacteria but unrelated to known RNase families. The cryo-EM structure of RNA bound to Escherichia coli YicC endonuclease suggested a clamshell-like ribonuclease mechanism for cleavage, although its exact mechanism and function remain elusive. Here, we report new cryo-EM structures that provide snapshots of the closing of the complex and also capture a surprising dimeric-RNA:protein complex. We further characterize the RNA cleavage targets in solution using nuclear magnetic resonance (NMR) spectroscopy, and use all-atom molecular dynamics (MD) simulations to study Mg^2+^ coordination. Our findings suggest that the YicC family can bind RNA hairpin structures and dimeric conformations with appropriate sequence and structural constraints, with cleavage driven strongly by Mg^2+^ ion localization. This study therefore provides overall insight into the novel cleavage mechanism of the highly conserved family of YicC-like endoribonucleases.

## Introduction

RNase enzymes have many essential functions in bacteria, including regulating mRNA levels to allow bacteria to respond to changes in the environment.^1, 2^ We were recently able to identify a new endonuclease, YloC, in a *B. subtilis* strain that lacks all four known 3’ exoribonucleases (PNPAse, RNase PH, RNase R, and YhaM).^3^ YloC belongs to the YicC superfamily of proteins, a highly conserved but previously uncharacterized family of RNases, found in both Gram-positive and Gram-negative bacteria. Remarkably, this enzyme family lacks sequence similarity to any known RNases, indicating a novel structural fold. Consistent with other RNase classes, this enzyme is divalent-cation dependent (Mg^2+^, Mn^2+^, and Co^2+^) and exhibits a putative preference for single-stranded RNA substrates.^3^ There have been reports of the effect of YicC family members on the processing of a small RNA in *E. coli* ^*4*^ and sporulation in *Clostridioides*^5, 6^, but beyond these known biochemical traits, the physiological substrates of the bacterial YicC family remain uncharacterized, highlighting a significant gap in our understanding of this protein group.

Recent crystallographic studies by our group and others have identified that the prototypical YicC member from *E. coli*, YicC, forms a hexamer.^7, 8^ The architecture consists of two trimers that adopt an expanded conformation anchored by a hexameric cap. We subsequently determined a cryo-EM structure of YicC bound to a synthetic 26-mer RNA oligonucleotide, called 2051. Notably, the binding of this single-stranded RNA induces a conformational shift: the two trimers collapse into a closed barrel that encapsulates the 2051 RNA, which itself adopts a hairpin fold. Within this asymmetric hexamer, we identified a putative active site featuring two possible sites for Mg^2+^ ions coordinated by Glu216 and Glu217. Although Mg^2+^ was not present in the sample, we observed a discrete density that we assigned to water molecules mimicking the coordination effects of Mg^2+^ ions. The coordination of Mg^2+^ ions by a glutamate pair is a common feature of the active site of divalent cation dependent-RNases,^9^ suggesting a case of convergent evolution.

Characterization of the catalytic activity of YicC revealed that the 26-nt 2051 RNA is cleaved at two distinct sites. The primary cleavage event (Site 1) produces a major 23-mer product (called 2182) and aligns precisely with the Mg^2+^ coordination site identified in our cryo-EM structure, suggesting a catalytically competent conformation. Conversely, the secondary cleavage at Site 2 suggested either RNA translocation or the presence of multiple active sites. We proposed a model wherein the substrate adopts heterogeneous hairpins in solution, one with a hairpin near the 5ʹ end and another with a hairpin near the 3ʹ end. Preferential cleavage at the 5ʹ end of these distinct hairpins accounts for both observed fragments. Lastly, we found a catalytically inactive variant (E281A) that exhibits a ten-fold increase in binding affinity compared to the wild-type protein (likely due to the decrease in negative charge from the mutation in all six monomers), providing a robust tool for capturing stable enzyme-substrate complexes.

In this study, we use cryo-EM and biophysical approaches to elucidate the structure and sequence requirements for YicC-mediated RNA processing. We demonstrate that substrates incapable of forming the canonical 5ʹ hairpin instead adopt non-canonical duplexes that are stabilized by both Watson-Crick-Franklin and Hoogsteen base-pairing within the positively charged protein cavity. Integration of structural data with nuclear magnetic resonance (NMR) spectroscopy suggests that these specific RNA folds are protein-induced, forming only upon sequestration within the YicC barrel. These findings provide new insights into the RNA-protein recognition and unique catalytic mechanism of this newly discovered RNase family.

## Results

### Alteration of the cleavage site leads to a novel substrate for the enzyme with a single cleavage event

We previously identified a synthetic YicC RNA substrate, 2051 (Table S1), which undergoes efficient cleavage at two distinct sites: 3ʹ to C3 (yielding fragments B and F) and to U14 (yielding fragments C and E). We showed that the F fragment (2182) is itself a good substrate for YicC, which generates two fragments by cleavage at Site 2 (D and E). A fifth fragment (D) also arises due to the potentially simultaneous cleavage of the full-length artificial RNA (Fig. 1A).^7^ To characterize the mechanism for cleavage at the second site (U14), we performed a cleavage reaction using a 5ʹ fluorescently labeled 2051 oligo under single turnover conditions and monitored the reaction over time. Two cleavage products are visible from the earliest time points, fragments B and C, suggesting that cleavage at Site 2 can happen independently of Site 1 binding and cleavage (Fig. 1B). Indeed, both the cleavage kinetics for fragments B and C follow pseudo-first-order behavior, reaching plateaus of 0.433 and 0.225, respectively. The calculated rate constants (K_obs_) for both fragments were similar (0.128 s^-1^ and 0.060 s^-1^ for fragments B and C), suggesting that while the final yields differ, the catalytic events leading to these fragments occur on a similar time scale (Fig. 1C).

**Figure 1.**
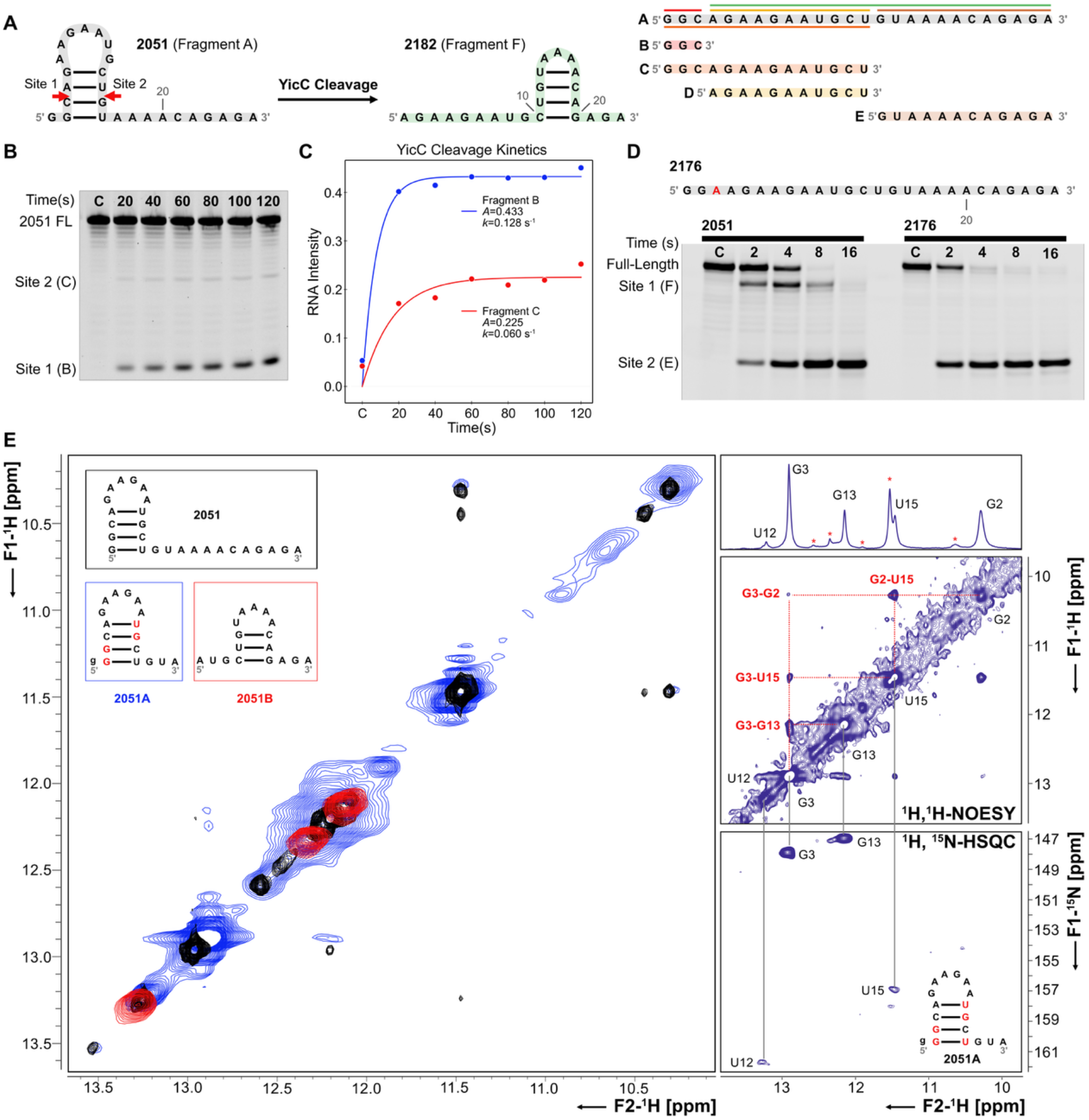
Cleavage and conformation of RNA fragments. (A) Sequence of 2051 and its five cleavage products, fragments B-F. Cleavage fragments are colored based on sites of cleavage (B) Single-turnover cleavage of 5ʹ labeled 2051 analyzed by gel electrophoresis. Reaction was terminated at the indicated times, or control (C) with no enzyme. (C) Cleavage kinetics from single-turnover experiment showing band intensity of cleavage products from site 1 and site 2 cleavage. (D) Sequence of the 2176 mutant transcript with cleavage assay with no Site 1 cleavage observed. (E) 2D ^1^H,^1^H NOESY overlays showing differences in solution structure between FL-2051, 2051A, and 2051B. 1D-^1^H spectra for 2051A is shown on the right, with the corresponding labelled ^1^H-^1^H NOESY and ^1^H-^15^N HSQC spectra shown directly below. Colored bases indicate base-paired imino positions with confidence. An additional G was incorporated into the design of 2051A to improve transcription efficiency with T7 polymerase.

To investigate the role of the 5ʹ hairpin, we designed 2176, a variant of 2051 in which C3 was mutated to A3, effectively disrupting 5ʹ-proximal hairpin formation (Fig. 1D).^10^ Using 3ʹ-labeled RNA, we observed that 2176 yields only fragment E upon completion of the reaction. In contrast, 2051 generates both fragments E and F, with the latter serving as a transient intermediate that is further processed into the final products D (lacking a 3ʹ label and thus not visible) and E. These results demonstrate that while the 5ʹ hairpin is strictly required for cleavage at Site 1, it is dispensable for cleavage at Site 2. Consequently, the dual-site cleavage of 2051 does not occur via simultaneous processing on both sides of a single hairpin, thereby ruling out a concerted cleavage model.

To validate the structural ensemble representing the 2051 hairpin in solution, along with identifying whether 2182 forms the 3ʹ hairpin configuration, we carried out both 1D-^1^H NMR and 2D-^1^H,^1^H NOESY (Fig. 1E, Fig. S1). In addition to the full-length sequences, we also designed truncated versions of the 2051 transcript that contained either the 5ʹ-proximal region (2051A) or the 3ʹ-proximal region (2051B) (Table S1). For homonuclear NMR experiments, these samples were purchased from IDT and resuspended in buffer (5 mM MgCl_2_, 7 mM NaCl, 50 mM KCl, and 50 mM NaH_2_PO_4_ buffer at pH 7.5, supplemented with 10% D_2_O), with a final concentration of 150 μM for each condition. From the 2D NOESY measurements, we observe a well-structured FL transcript with clear NOE crosspeaks, representative of through-space magnetization transfer of exchangeable protons. The 2051A proximal hairpin overlapped with the FL transcript to a high degree, with shared NOEs seen both upfield (∼10.4 ppm) and further downfield (12.3 ppm), and shared imino resonances across the diagonal.

Notably, stark differences were seen between the 2051A and 2051B hairpins, with some degree of overlap still observed between the 2051B and FL transcripts. These results reveal that both the 5ʹ and 3ʹ hairpin structures are sampled by the FL transcript in solution. In addition, the presence of distinct imino resonances for the FL transcript (10.5 ppm) suggests the sampling of alternative structured states that are not described by either 2051A or 2051B. To establish the secondary structure of the dominant 5ʹ-hairpin with greater detail, we turned to ^1^H, ^15^N-HSQC measurements. We transcribed a ^13^C, ^15^N-labelled variant of 2051A for improved resolution during structural analysis, resuspended in the same buffer. Evaluation of NOE crosspeaks and HSQC resonances suggests a 4 base-pair hairpin with a 6-nucleotide apical loop. Other peaks also exist in the 1D and NOESY spectra, which could correspond to alternative structures in solution or possible dimeric configurations. Overall, these findings support the hypothesis that the 2051 hairpin may exist in a diverse ensemble dominated by 2051A. Upon cleavage, this transcript could then shift to adopt a 3ʹ-dominant structure that is favored by 2182. This also supports the results of the cleavage assays described above, where fragments B, D, and E are not predicted to form structures in solution.

### Cryo-EM structure of 2176 bound to YicC

To elucidate the mechanism of cleavage at the second site, we performed cryo-EM experiments with the 2176 transcript in the presence of wild-type YicC. As described above, 2176 is cleaved *exclusively* at the second site. Surprisingly, we obtained several unique classes from the same sample with moderately high resolution. Solving these structures yielded three unique protein snapshots that span the trajectory from the open, apo structure to the closed RNA-bound structure (Fig. 2A and Fig. S2). Increasing amounts of RNA density in the core of the cavity were observed for each class, including the semi-closed state (Fig. 2B) and closed (Fig. 2C) state with the clearest density. We were able to fit the YicC backbone into the density for each structure, with enough resolution to detect conformational changes in the alpha-helical backbone. While the resolution was not high enough to capture atomistic density of the RNA itself, fitting 2051 onto the density map showed density in the binding site that could not be accounted for by the 2051 hairpin alone (Fig. 2D). The series of structures from this data set provides snapshots of a possible reaction pathway between the apo and closed states, highlighting the dynamic nature of the protein and how the RNA can aid in building the catalytic site from disparate monomers of the protein within the closed conformation. However, the unaccounted density in the structure shows that 2176 does not bind with the same mechanism as 2051.

**Figure 2.**
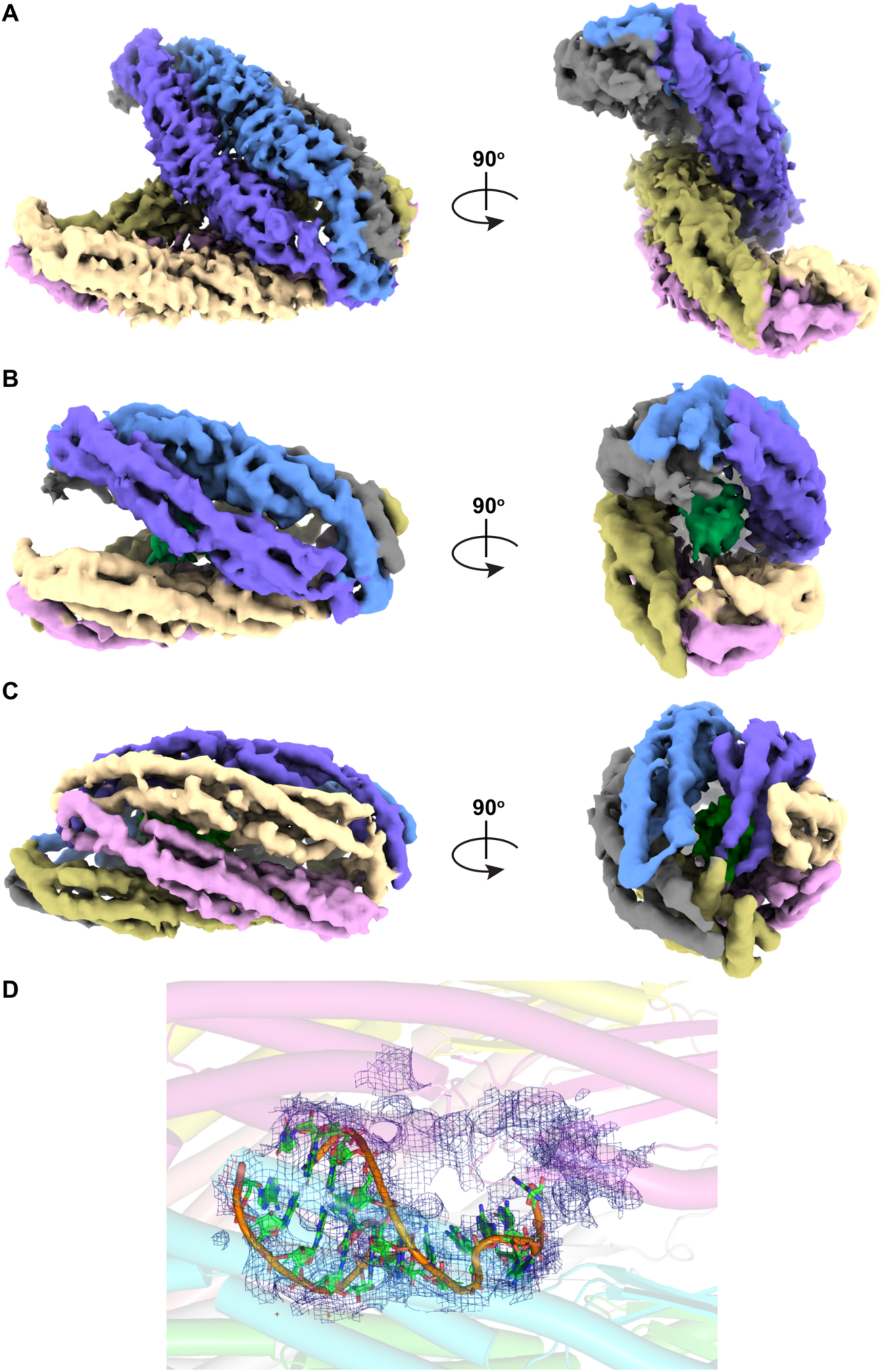
Structure of YicC bound to 2176 RNA in different conformations from the same sample. Density is colored according to chain, with the RNA colored in dark green. Two different angles are shown for each structure. (A) open conformation. (B) partially closed conformation. (C) fully closed conformation. (D) close-up view of fully-closed conformation with non-protein electron density shown in blue. The model shown is 2051 bound to YicC (PDB 8VES), with RNA depicted in green and orange with corresponding density in blue.

### Cryo-EM structure of 2182 bound to mutant protein

In order to obtain a higher-resolution structure, we turned to the E281A mutant of YicC, which binds RNA with ten-fold higher affinity but does not cleave RNA, likely due to the decrease in negative charge from the mutation in all six monomers. We incubated E281A YicC with 2182 RNA and analyzed the samples by cryo-EM. After processing cryo-EM data, we observed that a majority of the complexes (73.8%) were in the closed conformation, likely due to the increased affinity of the mutant for RNA substrates (Fig. S3). After solving the structure (Table S2), we saw that the protein conformation is nearly identical to the previous structure with 2051. However, when we built the RNA sequence, much to our surprise, instead of observing the 3ʹ hairpin that 2182 forms in solution, we observed the 2182 RNA binding as a dimer in the active site (Fig. 3A). The protein forms a closed barrel that is a pseudo-dimer of trimers that encloses the RNA substrate (Fig. 3B), as seen in the previous structure.^7^ The RNA sits inside the barrel of the YicC hexamer with the helical axis parallel to the YicC long axis. If we look inside the barrel, we can observe the two strands wrapping around each other. As in the previous structure, many of the contacts between the RNA and protein are driven by electrostatics, mainly of arginine side chains (Arg 30/E, Arg 30/D, Arg 280/B, Arg 280/A, Arg 30/A) that interact with the negatively charged phosphate backbone of each RNA strand (Fig. 3C). In addition, there are numerous other contacts to the ribose moieties: Arg 251/A, Thr 255/A, Asn 250/C. The duplex consists of both Waston-Crick-Franklin and Hoogsteen base pairing extending across the entire duplex, with the exception of the adenine in the middle of the duplex that is extruded on both sides. The dimer also explains the extra density in the wild-type structure (Fig. 2D), showing that the wild-type protein likely binds a 2176 dimer as the major species.

**Figure 3.**
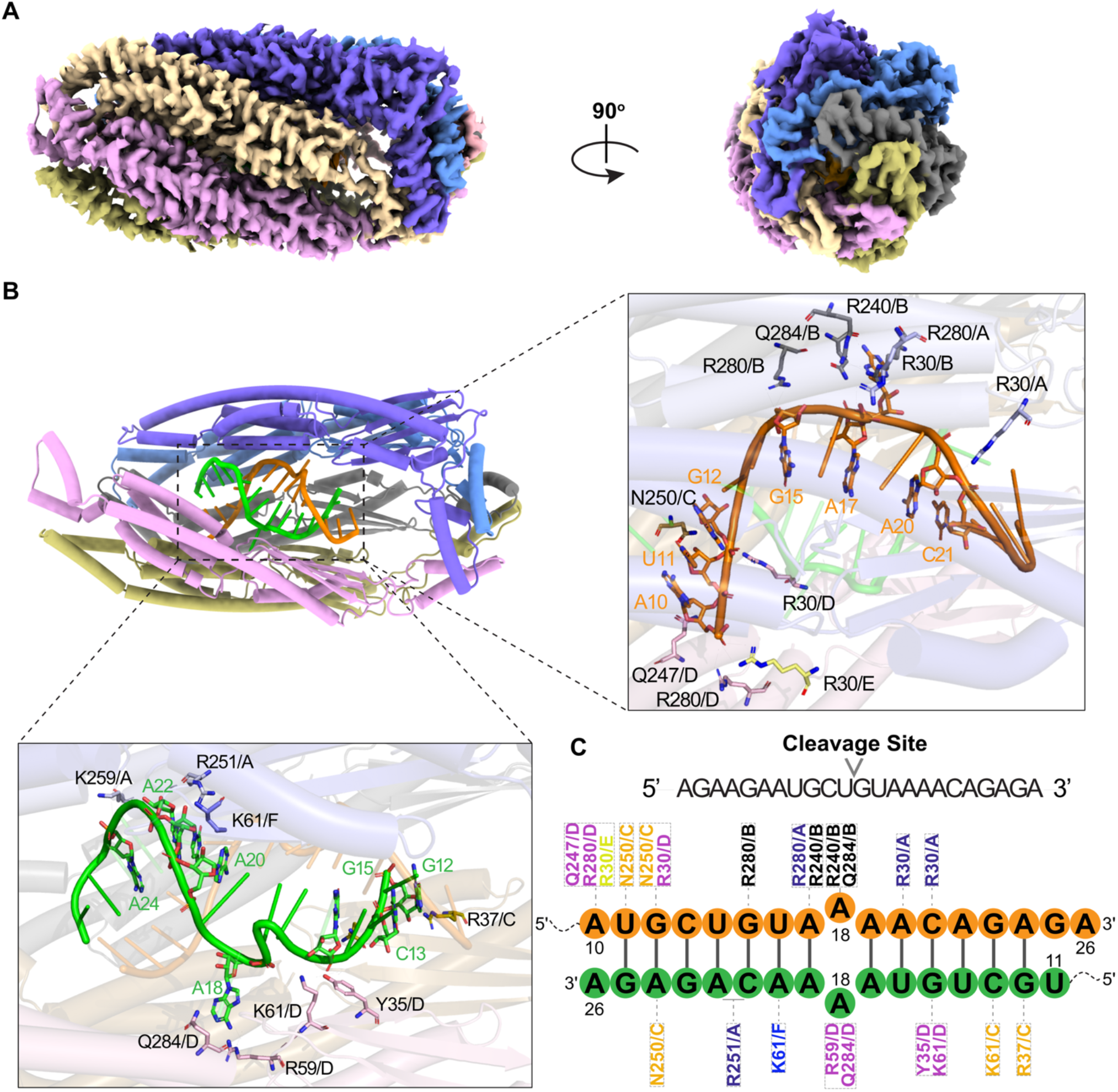
Structure of YicC E281A bound to 2182 RNA. (A) Overview of protein-domain architecture highlighting hexamer barrel conformation. (B) Overall structure of E281A:2182, shown in cartoon form, colored by chain. RNA dimer is colored by strand and is visible inside the cavity. Both RNA strands in the dimer are shown (orange, green) along with corresponding protein residues coordinating with backbone positions. (C) Schematic of observed interactions between YicC and the RNA dimer, with residue and chain indicated for each amino acid.

When we overlay our previous structure (PDB 8VES, wild-type protein bound to 2051) with this second high-resolution RNA complex structure, we find that many features are highly conserved (Fig. S4). The RNA backbone binds nearly identically for the two complexes. In addition, the extruded adenine base from one strand overlaps exactly with the 2051 structure and fits into a pocket formed by Arg 240 on one strand and Arg 59 on the other. This suggests the adenine pocket is a conserved feature of these substrates and might provide some sequence specificity.

Because we were surprised to see the RNA bound as a dimer, we were concerned that it might be an artifact of the mutant YicC protein. Therefore, we obtained an additional cryo-EM structure. We used E281A YicC in a complex with 2051 to see if the mutant causes dimerization. However, the E281A:2051 structure (Fig. S5-6) is identical to the previous wild-type:2051 structure, suggesting that the mutation does not induce dimerization of RNA. We also wanted to see if 2182 RNA can dimerize in solution, since it was predicted to be a hairpin and NMR was consistent with the hairpin structure. Using a gel assay, we found that 2182 does not dimerize in solution, but fully complementary control RNA oligonucleotides do (1935:1955; Fig. S7), showing that the dimerization of 2182 requires the protein.

### Catalysis of the dimer

We then examined the structure to understand how cleavage could occur on this dimer. The YicC hexamer forms an asymmetric pseudodimer featuring a proposed single, functional catalytic site consisting of a triad of glutamates (Glu216, Glu217, Glu281). In the E281A:2182 structure, the RNA cleavage site (between bases 14 and 15) is not adjacent to this triad, including the 216/217 from chain A that are the catalytic sites in the previous 2051 structure (Fig. 4A). To gain insight regarding the properties of the cleavage site, we used all-atom MD simulations. Existing literature on bacterial endo- and exonucleases describes a catalytic mechanism that is dependent on divalent cations within the active site. In the 2051 structure, a density was observed that could correspond to a divalent cation such as Mg^2+^. We simulated both the 2051 and 2182 transcripts bound to WT YicC protein in solution for 1 μs in triplicate. Ion mapping and radial distribution function (RDF) calculations were performed to visualize and quantify localization (Fig. 4B, Fig. S8). For the WT protein bound to 2051 (PDB ID: 8VES), we see repeated localization of at least two magnesium ions to E216/A, E217/A, and E281/F. Both ions were observed populating the cleavage site between positions C3 and A4, positioned adjacent to the backbone. To quantify this distribution, we calculated the RDF in a pairwise manner to describe the specificity of Mg^2+^ to this cleavage site and compared this to Na^+^ localization. The RDF was calculated for all Mg^2+^ atoms within a 7 Å cutoff radius of RNA nucleotides at this site (C3, A4, G5, A6). We observed that Mg^2+^ showed high distribution probabilities around these positions ranging from 2 to 6 Å. The same calculations for Na^+^ showed little to no preferential distribution at the cleavage site. This suggests an Mg^2+^-dependent cleavage of the scissile phosphate group between C3 and A4, where the cations are potentially further stabilized within the catalytic glutamate triad (E216, E217, E281). We also performed MD simulations with identical conditions for the E281A variant bound to 2182. Here, ion mapping calculations showed a lone Mg^2+^ that localized to the cleavage site. Notably, the loss of a second divalent cation at this position potentially explains why the mutant is inactive, most likely because the glutamate substitution (Fig. S9) prevents the second Mg^2+^ ion from binding.

**Figure 4.**
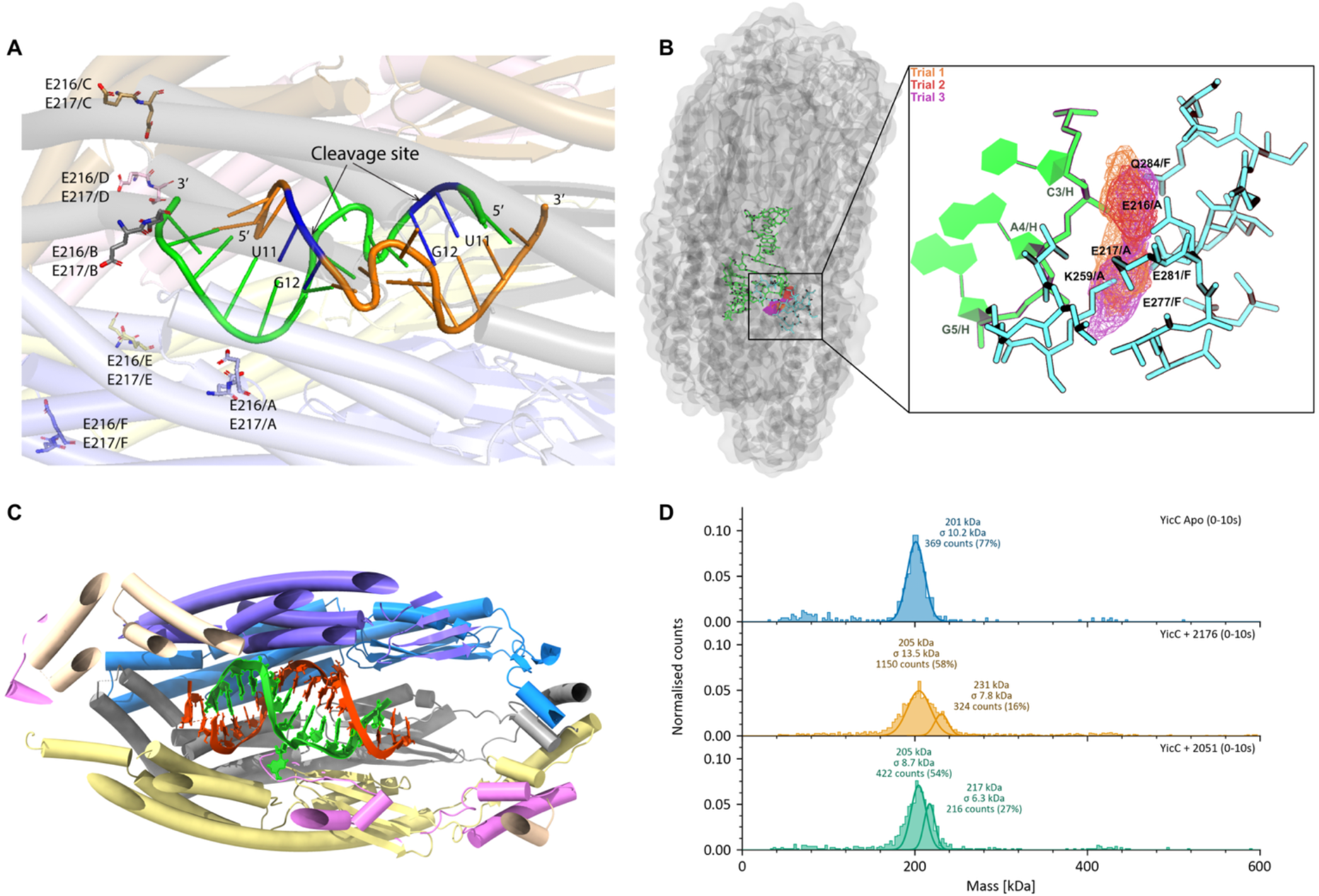
Catalytic site of YicC. (A) Closeup perspective of E281A:2182 structure with cleavage site indicated. Catalytic glutamates are also highlighted flanking the RNA with residues and chains identified. (B) Molecular dynamics simulations of WT YicC bound to 2051A. Ion density mapping for Mg^2+^ is shown for all three trials and is depicted by orange, red, and purple wireframes. (C) Overall structure of E281A:2176 with Mg^2+^. Mutant YicC is colored by chain and the RNA dimer is shown as a filled cartoon. (d) Mass Photometry showing mass of mutant YicC alone, bound to 2176, or bound to 2051, as indicated. The different molecular weight peaks are indicated.

Given the importance of magnesium for RNA folding,^11^ we wondered whether the dimer was formed due to the lack of magnesium. Revisiting 1D-^1^H NMR experiments, we compared the folding of the FL-2051 hairpin either in the presence or absence of 1.5 mM Mg^2+^ in a low-salt buffer (15 mM NaH_2_PO_4_ pH 7, 25 mM NaCl, 10% D_2_O). Upon addition of MgCl_2_, we see that while some upfield peaks are absent, the shift pattern remains largely the same regardless of Mg^2+^ being present (Fig. S10). Another possibility is that the structure of the RNA bound to the protein is dependent on Mg^2+^ and that the dimer is an artifact of RNA binding without Mg^2+^. Therefore, we resolved another cryo-EM structure of the 2176 substrate bound to E281A YicC in the presence of 5 mM MgCl_2_. We found that under these conditions, the RNA still forms a dimer bound to YicC (Fig. 4C, Fig. S11). Because the RNA is longer, we observe more RNA density relative to the 2182 RNA-YicC structure; nevertheless, the dimer structure is otherwise identical to the 2182 structure. Therefore, we conclude that magnesium does not help the RNA form a different structure when complexed with the YicC protein.

### RNA dimer formation in the protein in solution

As previously described, dimerization of the RNA is likely induced by protein binding. To further confirm this, we used mass photometry to look at binding of RNA to protein in solution. This would exclude the possibility that dimer formation is a function of the cryo-EM conditions or high protein concentrations. Using the E281A mutant protein, we examined the mass of protein alone or bound to 2176 or 2051 (Fig. 4D). Although the resolution is right at the limit for seeing 26-base oligonucleotides, it is clear that 2176 shows a peak with a substantially higher increase in mass than 2051, which is consistent with two copies of 2176 binding but only one copy of 2051 binding to the YicC hexamer.

### Cleavage and binding requirements

Lastly, we wanted to examine the binding and cleavage requirements of YicC. We initially reported that YicC had a requirement of single-stranded RNA and that double-stranded RNA was not a substrate.^3^ When we later solved the structure of 2051 bound to YicC, it became clear that hairpins are appropriate substrates.^7^ With the discovery of the RNA dimer structure, we wondered again whether duplex RNA could bind or be cleaved. Therefore, we designed RNA duplexes consisting of either perfectly complementary strands (2051:2292) or complementary strands with a mismatch in the middle (2051:2305) to increase the intrinsic dynamics and accessibility of the cleavage site. The RNAs were then annealed prior to cleavage or binding experiments (Fig. S12). Neither duplex was cut by YicC. However, fluorescence anisotropy experiments showed that both wild-type and mutant proteins can bind strongly to the dimers (30 μM and 4 μM, respectively, Fig. S13). YicC can bind double-stranded RNA but cannot cleave it. Our mismatch RNA was not cleaved, indicating that the rules for substrates are more nuanced. One surprising result was that although 2051 RNA binds much more tightly to the E281A protein than to WT-protein as we previously reported, the duplex RNAs bound less strongly to the mutant protein than the single-stranded RNA, indicating that some binding differences between the mutant and WT proteins remain.

## Discussion

In the short time since the discovery of YicC endoribonuclease activity, the exact cleavage mechanism remains elusive despite the structural features being increasingly described. Recent efforts have identified the involvement of other YicC family members in the processing and cleavage of small RNA in *E*.*coli*^12^ and sporulation in *Clostridioides*^13^, but the mechanisms have not been thoroughly established. Here, we report several new cryo-EM structures of YicC bound to different RNA substrates to try to better establish the cleavage mechanism. These structures, accompanied by cleavage assays and biophysical measurements, aim to answer the elusive question: how does an RNA oligonucleotide get cleaved at two precise positions by YicC RNases that are not sequence-specific? The cryo-EM structures show that the RNA substrate binds dimerically to the protein, making the proclivities of the enzyme appear less trivial than initially presumed.

We turned to NMR to understand the structures of the RNA transcripts in solution. While 2051 forms a 5ʹ hairpin in solution, it is different from the 5ʹ hairpin we observe bound to the protein. Substrates that cannot form the 5ʹ hairpin, like 2182, form a 3ʹ hairpin in solution but form a dimer bound to protein. YicC is unique among RNases in that it binds a single substrate in a hexameric protein via a clamshell mechanism. Here, binding is driven by a series of arginines that decorate the inside of the barrel and bind to the negatively charged RNA phosphate backbone. Therefore, it is perhaps not surprising in hindsight that the protein plays such a significant role in the folding of the RNA, given that much of the binding energy comes from the protein-RNA complex rather than the RNA folding alone. Protein-induced conformational changes of RNA are quite common in the literature.^12-14^ This makes it especially challenging to predict the fold of an RNA that is bound to this protein. While the 2182 and 2176 RNA oligonucleotides can form weakly stabilized hairpins in solution, when bound to the protein, they form a variety of uncommon interactions, including G(anti)-G(syn) and G(anti)-A(syn) Hoogsteen pairs that are driven by the positively charged interior of the barrel, compelling the RNA to adopt a longer dimeric structure.^15, 16^

We previously solved only a single structure of RNA bound to the YicC family, making it challenging to determine the required features needed to form a complex. Now, with additional structures, we can make further observations. While the protein can bind double-stranded RNA quite well, all the substrates have single stranded components or a mismatch that enables some degree of flexibility in the structure of the RNA, perhaps acting as a prerequisite for cleavage. We also note a conserved extruded adenine base seen in all structures.

With the current set of experiments on the YicC cleavage process, we propose a model (Fig. 5), where YicC binds an RNA substrate like 2051 and induces a fold that consists of a 5ʹ hairpin (Fig. 5B). This fold is highly favored in the context of the protein and so cleavage occurs at the base of this hairpin to generate the first cleavage event that is preferred (Fig. 5D). The larger fragment of this cleavage, 2182 RNA, preferably forms a dimer bound to the protein, and this dimer is then cleaved (Fig. 5E). The 2051 substrate can also be cut at site 2 initially, as seen in our single-turnover experiments. We hypothesize that 2051 can bind the enzyme as a dimer (Fig. 5A) or a 3ʹ-hairpin (Fig. 5C), which is what allows cleavage at site 2. We note that in the 2182 dimer structure, the cleavage site is not near the active site. There are several possible explanations for this. First, it is possible that the observed dimer is a pre-cleavage binding state or transitory intermediate that is required for cleavage after an additional, not yet identified, conformation is sampled. Second, since the resolution was not high enough in the WT protein to accurately place the dimer, we cannot exclude that this form of the dimer is different from that which binds to the mutant E281A protein variant. Third, we also cannot rule out the possibility that the 3ʹ hairpin of 2182 is the active cleavage structure and the dimer is an inactive substrate of the enzyme. If the 3ʹ hairpin is not favored but can still be cleaved efficiently, it might be difficult to observe when compared to an inactive dimer. Future work will be required to narrow down these possibilities.

**Figure 5.**
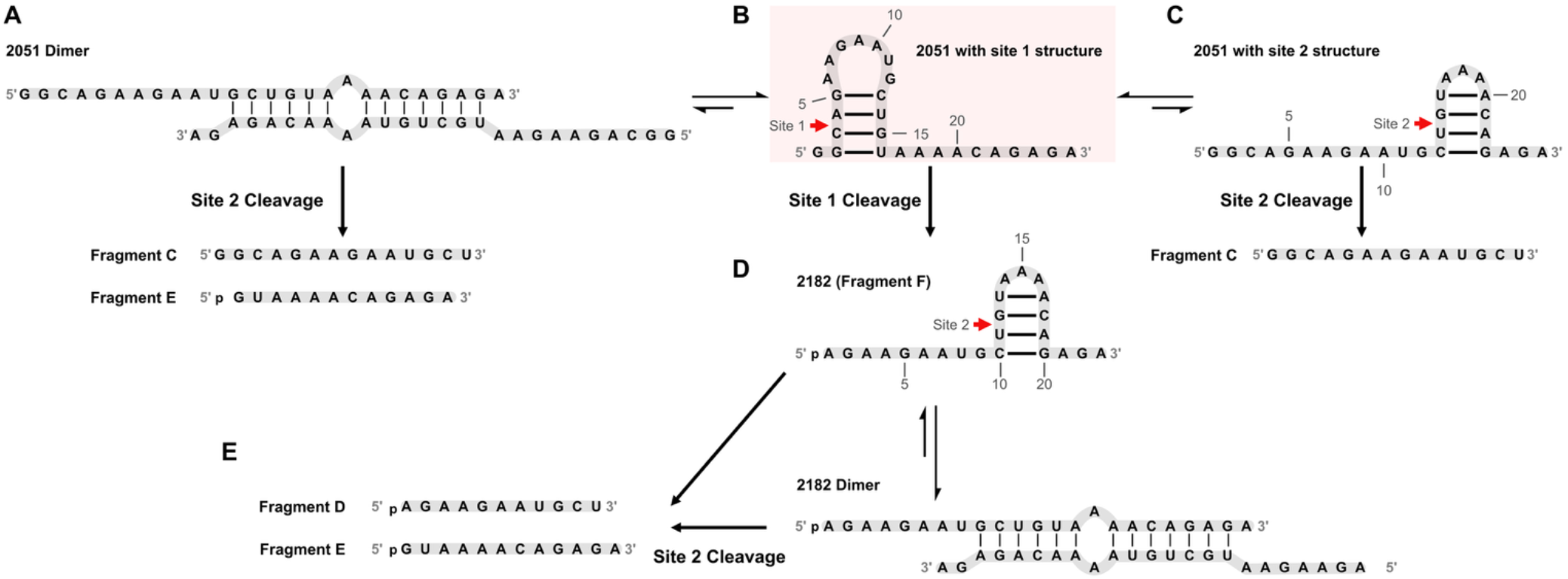
Proposed model for two cleavage events on 2051 RNA substrate. (A) Proposed 2051 dimer which yields fragments C and E upon cleavage at site 2. (B) Preferred structure of 2051 with the 5ʹ-proximal hairpin and cleavage at site 1. (C) Possible alternate conformation of 2051 with cleavage at site 2, yielding fragment C. (D) Site 1 cleavage product of 2051 yields fragment F (2182), which samples both a 3ʹ-proximal hairpin conformation and adimer. (E) Site 2 cleavage fragments of 2182, yielding fragments D and E.

Finally, the new structures also show that, because of the ability to induce RNA dimerization, it is possible that this family of enzymes is used to cleave RNA duplexes that do not come together on their own, in addition to hairpin structures. Identification of physiological substrates of the YicC family will confirm whether hairpins or duplexes, or both, are substrates.

## Materials and Methods

### Protein purification

YicC WT and YicC E281A were previously cloned into His-sumo expression vectors ^7^ and expressed in *E. coli* BL21 LOBSTR (Kerafast) expression cells. Cultures were grown at 37 °C to an OD_600_ of approximately 2.0, then cooled to 30°C and induced with 2 mM isopropyl β-D-1-thiogalactopyranoside (IPTG) in the presence of 2% glycerol. After 5 hours of expression, cells were harvested by centrifugation and resuspended in lysis buffer (TBS containing 20 mM Tris-HCl and 250 mM NaCl). Cells were lysed by sonication and the lysate was clarified by centrifugation. The supernatant was applied to Ni-NTA resin pre-equilibrated with binding buffer (20 mM Tris-HCl, 250 mM NaCl, 40 mM imidazole). The resin was washed with wash buffer (20 mM Tris-HCl, 250 mM NaCl), and protein was eluted with elution buffer (20 mM Tris-HCl, 150 mM NaCl, 250 mM imidazole, 10% glycerol). The eluted protein was treated with SUMO protease for tag cleavage and analyzed by SDS-PAGE. The sample was further purified by ion-exchange chromatography using a HiTrap Q column and then finally on a Superdex200 increase size exclusion column.

### RNA labeling and RNase assay

RNA oligonucleotides were fluorescently labeled at the 5ʹ or 3ʹ ends as previously reported.^17^ The RNase assay was performed using 5ʹ or 3ʹ labeled oligonucleotides and the corresponding unlabeled nucleotide in 5-fold excess, incubated at 37ºC for increasing time points. Reaction products were run on a 15% or 20% polyacrylamide denaturing gel and imaged on a Li-COR Odyssey. For single turnover assay, the reaction used a 12:1 ratio of enzyme to RNA. For other cleavage assays a ratio of 1:20 was used.

### In-Vitro Transcription

Unlabelled RNA oligonucleotides (2051, 2051A, 2051B, 2182, 2182A, 2182B) for 1D 1H imino and 2D NOESY experiments were purchased from IDT. Isotopically labelled ^13^C-^15^N labelled RNA (2051A) was prepared by in-vitro transcription with in-house T7 RNA polymerase using DNA oligonucleotide templates also purchased from IDT, with labelled rNTPs purchased from Silantes. The RNA was purified using HPLC, followed by desalting and lyophilization. Samples were resuspended in the chosen NMR buffer and refolded by heating at 95 °C and snap-cooling on ice. Concentrations were measured using a NanoDrop Spectrometer.

### Electron microscopy

YicC-RNA complex sample at a concentration of 5 mg/ml was supplemented with 0.05% dodecyl maltoside detergent (DDM) immediately before plunge-freezing to enable even distribution of particles on the grid. 3 μl of sample was applied to glow-discharged Quantifoil holey carbon grids (Cu, R1.2/1.3, 300 mesh). The grids were blotted for 2 s and plunged into liquid ethane with a Vitrobot plunger (4°C and 90% humidity). Cryo-EM data were collected with a Titan Krios microscope (FEI) operated at 300 kV. For YicCE281A/RNA 2182, movies were collected using Leginon^18^ at 105,000X nominal magnification with the calibrated pixel size of 0.4130 Å. The dose rate was 33.77 e^-^/Å^2^/s with a total exposure of 1.60 seconds, for an accumulated dose of 54.03 e^-^/Å^2^. Intermediate frames were recorded every 0.04 seconds for a total of 40 frames per micrograph. A total of 13433 images were collected at a nominal defocus range of -0.2 – 2.7 μm. For YicCE281A/RNA 2051, movies were collected using Leginon at 81,000X nominal magnification with the calibrated pixel size of 0.4280 Å. The dose rate was 32.19 e^-^/Å^2^/s with a total exposure of 1.60 seconds, for an accumulated dose of 51.50 e^-^/Å^2^. Intermediate frames were recorded every 0.04 seconds for a total of 40 frames per micrograph. A total of 14384 images were collected at a nominal defocus range of 0.8 – 2.0 μm. For YicCWT/RNA 2176, movies were collected using Leginon at 105,000X nominal magnification with the calibrated pixel size of 0.4128 Å. The dose rate was 25.89 e^-^/Å^2^/s with a total exposure of 2.0 seconds, for an accumulated dose of 51.77 e^-^/Å^2^. Intermediate frames were recorded every 0.04 seconds for a total of 50 frames per micrograph. A total of 14170 images were collected at a nominal defocus range of -0.3 – 2.9 μm. The statistics for the cryo-EM data are listed in the Supplementary Table. For the YicC E281A/RNA 2176 structure, 3 μL of E281A YicC at a concentration of 2.7 mg/mL, incubated with RNA fragment 2176 in a 2:1 ratio, 5 mM MgCl_2,_ and 0.05% DDM, was applied to glow-discharged Quantifoil holey carbon grids (Cu, R1.2/1.3, 300 mesh). Grids were blotted for 3 seconds and plunge-frozen in liquid ethane using a Vitrobot (4°C, 90% relative humidity). Data were collected on an EF-Krios transmission electron microscope operated at 300 kV and equipped with a Gatan K3 direct electron detector at a nominal magnification of 81,000×, corresponding to a calibrated pixel size of 0.85 Å.^18^ The detector was operated in super-resolution mode at the end of a GIF-Quantum energy filter with a slit width of 20 eV. Movies were recorded at a dose rate of 41.14 e−/Å^2^/s for a total exposure time of 1.36 s, resulting in an accumulated dose of 55.95 e−/Å^2^. Intermediate frames were saved every 0.03 s, yielding 40 frames per micrograph. In total, 9697 movies were collected at a nominal defocus range of 0.6–1.6 μm.^19^

### Image Processing and Model Building

Movies were motion-corrected using MotionCor2^20^ and imported to cryoSPARC^21^ for further processing. Contrast transfer functions were estimated using patchCTF in cryoSPARC. An initial model was produced from a subset of micrographs using blob picking, followed by extraction, 2D classification, selection of key classes, and generation of a model ab initio. Subsequent map models were produced from a curated micrograph set using particles found by picking using the initial map model as a template. Particles were extracted, subjected to 2D classification, and a final particle stack was obtained by iterative rounds of 3D classification, generating several bad models from rejected particles as a sink in hetero-refinement. A final map was obtained using NU-refinement58. Specific processing procedures regarding YicC E281A/RNA 2182, YicCE281A/RNA 2051 and YicCWT/RNA 2176 can be found in Supplementary figures. A total of 11,099 dose-weighted micrographs of the YicC complex with RNA fragment 2176 were collected and imported. Processing was conducted using cryoSPARC^21^. CTF estimation for each micrograph was performed using Patch CTF. 3,552,024 particles were selected through blob picking and extracted from the micrographs. Ab initio models served as initial references for subsequent 3D classifications. The particles underwent multiple rounds of 2D and 3D classification to discard poor-quality particles, resulting in a final set of 166,251 cleaned particles used for NU-refinement^22^ with C1 symmetry, achieving a final map at 3.04 Å resolution. Structures were built by docking our previous structure (8VES) using Phenix^23^ and manually in Coot^24^, and refined using PHENIX with Phenix real_space_refine, incorporating B-factor refinement and rotamer corrections. Final maps were imported to PyMOL^25^ and UCSF Chimera^26^ to generate the shown in the manuscript.

### Mass photometry

YicC E281A at 2.5mg/ml was incubated with RNA2176 or R2051 on ice for 30 mins. Then, mass photometry tests were conducted using a Refeyn TwoMP. For data acquisition, we used AcquireMP (version 2025R2). All measurements were taken using MassGlass UC slides and cassettes (Refeyn MP-CON-21022), and in the regular image size and normal measurement mode. Calibration of the data was achieved using beta-amylase (Sigma A8781). 20 μl of room-temperature buffer was dispensed into a designated well, and the focal location was identified automatically. 0.1 μl of the YicC or YicC/RNA complex with a final concentration post-dilution of around 62.5 nM was incorporated into the well and mixed rapidly in order to obtain signal before the complex dissociated. Mass photometry readings spanned 60 seconds, but data was analyzed in the first 10 seconds before the complex dissociated due to the dilution. Analysis was conducted and figures were generated through the DiscoverMP software (version 2025R2).

### Fluorescent anisotropy (FA)

FA experiments were performed mostly as previously described.^7^ RNA 2051 was chemically labeled at the 3’ end with FTSC and gel purified and then annealed with 2305, 2292, or nothing. After duplex formation was verified by native gel, protein that was diluted at the indicated concentrations was added in, and anisotropy measured in fluorescent polarization mode on a Victor Nivo plate reader with appropriate filters. Data was fit usingGraphPad Prism to determine binding constants.

### Molecular dynamics (MD) general protocols

Molecular dynamics (MD) simulations of the YicC endonuclease-RNA complex were carried out using the PMEMD module of Amber v23 software on NVIDIA L40S and V100 graphical processing units (GPUs)^26^. For RNA structures, the *LJbb* force field parameter set was used, which combines OL3 parameters with updated van der Waals parameters for phosphate oxygens adapted from Steinbrecher and Case^28,29^. The YicC endonuclease was modelled as a hexomer using the ff14SB parameter set to improve modelling of side-chain rotamers and backbone secondary structures relevant to RNA-protein interactions in solution^30^. The solvent was described with the TIP3P water model, and mono- and divalent ions were captured with the Li-Merz 12-6-4 ion parameters.^31^ Bonds involving hydrogen atoms were treated with the SHAKE algorithm with a 0.00001 Å tolerance for restraints. Long-range Coulombic interactions were calculated using the particle-mesh Ewald method using a 10 Å cutoff^32^. Production runs were carried out in the NPT ensemble using Langevin dynamics with a collision frequency of 1.0 ps-1. A Berendsen barostat was used to maintain a pressure of 1 atm at 298.15 K^33^. All production runs were performed with a 4.0 fs timestep using hydrogen mass repartitioning^34^. All trajectory processing and analysis were performed with the CPPTRAJ module of AmberTools, with visualization carried out with Visual Molecular Dynamics (VMD).^35, 36^

#### MD simulations of YicC endonuclease-RNA complexes

MD simulations of the YicC endonuclease were performed from Cryo-EM structures. These included the YicC endonuclease bound to a 26-nt RNA substrate (2051, PDBID 8VES), the same endonuclease bound to 2182, and the YicC protein in an apo state (PDBID 8VER). Hydrogen atoms were added as needed if absent from structure files. The systems were solvated in a 14 Å truncated octahedron with periodic boundary conditions. Neutralizing Na^+^ counter-ions and a 5 mM MgCl_2_ buffer were included using the tleap module of AmberTools2024. The final system sizes were 290,417 atoms for the 2051 complex, 290,690 atoms for the 2182 complex, and 294,622 atoms for the YicC WT protein alone.

The solvent was minimized for 1,000 steps following the steepest descent algorithm and then for 9,000 additional steps using conjugate gradient with positional restraints applied to the remainder of the system (1 kcal/mol*Å^2^). The systems were heated to 298.15 K with increased restraints (100 kcal/mol*Å^2^) on all atoms apart from hydrogens and water molecules. After equilibration with constant pressure (1 bar) for 1 ns, the restraints were decreased (10 kcal/mol*Å^2^) for an additional 1 ns of equilibration. Minimization was again carried out with 10 kcal/mol*Å^2^ positional restraints on RNA and protein backbone heavy atoms (N, α-C, C=O, C3, C4, C5, O3, O5, P) using 10,000 steps following conjugate gradient. After this minimization, a series of five-20 ns constant NPT equilibration simulations were performed with tapering restraints on the same backbone positions, where the force constant decreased in each step from 10 kcal/mol*Å^2^ to 5, to 2.5, to 1.0, to 0.5 kcal/mol*A2. All equilibration was performed with a 2 fs timestep. Production simulations were then obtained in triplicate for both systems for 1.2 µs with a 4 fs timestep and hydrogen mass repartitioning as described above. Analysis was performed on the last 1 µs of each simulation trial, yielding 9 µs of total processed MD simulation time.

### Nuclear Magnetic Resonance Spectroscopy

All NMR experiments were carried out using 150 μM unlabeled RNA for ^1^H-^1^D imino experiments with standard Bruker pulse programs on a Bruker Avance Neo 800 MHz spectrometer equipped with a TCI cryogenic probe^37, 38^. All experiments were performed in 5 mM MgCl_2_, 7 mM NaCl, 50 mM KCl, and 50 mM NaH_2_PO_4_ buffer at pH 7.5, supplemented with 10% D_2_O, and spectra were acquired at 278 K. The transcribed variant of 2051A was designed with an additional G nucleotide at the 5’-end to ensure successful transcription with in-house T7 polymerase. Structure prediction performed with the ViennaRNA packaged showed no changes in the predicted base-pair configurations in both the presence and absence of this additional nucleotide (Fig. S14). ^1^H-1D imino experiments were performed using 1024 scans, a total FID area of 4096, and a sweep width of 22.3 ppm. 2D ^1^H-^1^H NOESY experiments were optimized for mixing time by screening between 50 ms, 150 ms, and 300 ms, using 64 total scans and a FID area of 512 in the F1 dimension. For 2D ^1^H-^15^N HSQC experiments, 40 uM of ^13^C, ^15^N labelled 2051A transcript was evaluated using non-uniform sampling with a NUS amount of 50%. HSQC experiments were performed using 700 scans with a total FID area of 256 in the F1 dimension. All 1D and 2D spectra were processed and analyzed using Bruker TopSpin (4.4) and CCPN (3.3.2.3)^39^.

## Acknowledgements

We thank Kayleigh Fay for assistance with the mass photometry. Research reported in this publication was supported by National Institutes of Health award numbers GM147211 to D.H.B., R35GM124838 to M.B.L., and R35GM160278-0 to A.N.J. Some of this work was performed at the Simons Electron Microscopy Center at the New York Structural Biology Center, with major support from the Simons Foundation (SF349247). This work was supported in part through the computational and data resources and staff expertise provided by Scientific Computing and Data at the Icahn School of Medicine at Mount Sinai and supported by the Clinical and Translational Science Award (CTSA) grant UL1TR004419 from the National Center for Advancing Translational Sciences. MD simulations were conducted on the Torch High Performance Computer at New York University (NYU), supported by the NYU High Performance Computing resources, services, and staff expertise. NMR spectroscopy and CD experiments were carried out at the NYU Shared Instrumentation Facility, with support from Dr. Chin Lin and Dr. Joel Tang.

## References

1. Laalami S, Zig L, Putzer H. Initiation of mRNA decay in bacteria. Cell Mol Life Sci. 2014;71(10):1799–828. Epub 20130925. doi: 10.1007/s00018-013-1472-4. PubMed PMID: 24064983; PMCID: PMC3997798.

2. Hui MP, Foley PL, Belasco JG. Messenger RNA degradation in bacterial cells. Annu Rev Genet. 2014;48:537–59. Epub 20141001. doi: 10.1146/annurev-genet-120213-092340. PubMed PMID: 25292357; PMCID: PMC4431577.

3. Ingle S, Chhabra S, Chen J, Lazarus MB, Luo X, Bechhofer DH. Discovery and initial characterization of YloC, a novel endoribonuclease in Bacillus subtilis. RNA. 2022;28(2):227–38. Epub 20211123. doi: 10.1261/rna.078962.121. PubMed PMID: 34815358; PMCID: PMC8906540.

4. Chen J, To L, de Mets F, Luo X, Majdalani N, Tai CH, Gottesman S. A fluorescence-based genetic screen reveals diverse mechanisms silencing small RNA signaling in E. coli. Proc Natl Acad Sci U S A. 2021;118(27). doi: 10.1073/pnas.2106964118. PubMed PMID: 34210798; PMCID: PMC8271630.

5. Martins D, Salgueiro B, Sobral D, Gragera M, Hensel Z, Henriques AO, Romao CV, Serrano M. An endoribonuclease of the YicC-like family delays sporulation via sRNA degradation in Clostridioides difficile. Nucleic Acids Res. 2025;53(13). doi: 10.1093/nar/gkaf644. PubMed PMID: 40650975; PMCID: PMC12255296.

6. Martins D, DiCandia MA, Mendes AL, Wetzel D, McBride SM, Henriques AO, Serrano M. CD25890, a conserved protein that modulates sporulation initiation in Clostridioides difficile. Sci Rep. 2021;11(1):7887. Epub 20210412. doi: 10.1038/s41598-021-86878-9. PubMed PMID: 33846410; PMCID: PMC8041843.

7. Wu R, Ingle S, Barnes SA, Dahlin HR, Khamrui S, Xiang Y, Shi Y, Bechhofer DH, Lazarus MB. Structural insights into RNA cleavage by a novel family of bacterial RNases. Nucleic Acids Res. 2024;52(17):10705–16. doi: 10.1093/nar/gkae717. PubMed PMID: 39180400; PMCID: PMC11417398.

8. Huang L, Tam KS, Xie W. Structural and Biochemical Studies of the Novel Hexameric Endoribonuclease YicC. ACS Chem Biol. 2023;18(8):1738–47. Epub 20230803. doi: 10.1021/acschembio.3c00091. PubMed PMID: 37535940.

9. Yang W. Nucleases: diversity of structure, function and mechanism. Q Rev Biophys. 2011;44(1):1–93. Epub 20100921. doi: 10.1017/S0033583510000181. PubMed PMID: 20854710; PMCID: PMC6320257.

10. Barnes SA, Lazarus MB, Bechhofer DH. Cleavage specificity of E. coli YicC endoribonuclease. RNA. 2026. Epub 20260615. doi: 10.1261/rna.081057.126. PubMed PMID: 42297564.

11. Fürtig B, Richter C, Wöhnert J, Schwalbe H. NMR spectroscopy of RNA. Chembiochem. 2003 Oct 6;4(10):936–62. doi: 10.1002/cbic.200300700. PMID: 14523911.

12. Yamagami R, Sieg JP, Bevilacqua PC. Functional Roles of Chelated Magnesium Ions in RNA Folding and Function. Biochemistry. 2021;60(31):2374–86. Epub 20210728. doi: 10.1021/acs.biochem.1c00012. PubMed PMID: 34319696; PMCID: PMC8747768.

13. Hermann T, Patel DJ. Adaptive recognition by nucleic acid aptamers. Science. 2000;287(5454):820–5. doi: 10.1126/science.287.5454.820. PubMed PMID: 10657289.

14. Leulliot N, Varani G. Current topics in RNA-protein recognition: control of specificity and biological function through induced fit and conformational capture. Biochemistry. 2001;40(27):7947–56. doi: 10.1021/bi010680y. PubMed PMID: 11434763.

15. Williamson JR. Induced fit in RNA-protein recognition. Nat Struct Biol. 2000;7(10):834–7. doi: 10.1038/79575. PubMed PMID: 11017187.

16. Rypniewski W, Adamiak DA, Milecki J, Adamiak RW. Noncanonical G(syn)-G(anti) base pairs stabilized by sulphate anions in two X-ray structures of the (GUGGUCUGAUGAGGCC) RNA duplex. RNA. 2008;14(9):1845–51. Epub 20080724. doi: 10.1261/rna.1164308. PubMed PMID: 18658118; PMCID: PMC2525959.

17. Leontis NB, Westhof E. Geometric nomenclature and classification of RNA base pairs. RNA. 2001;7(4):499–512. doi: 10.1017/s1355838201002515. PubMed PMID: 11345429; PMCID: PMC1370104.

18. Barnes SA, Lazarus MB, Bechhofer DH. Cleavage specificity of E. coli YicC endoribonuclease. bioRxiv. 2026. Epub 20260326. doi: 10.64898/2026.03.25.714237. PubMed PMID: 41929057; PMCID: PMC13041888.

19. Suloway C, Pulokas J, Fellmann D, Cheng A, Guerra F, Quispe J, Stagg S, Potter CS, Carragher B. Automated molecular microscopy: the new Leginon system. J Struct Biol. 2005;151(1):41–60. doi: 10.1016/j.jsb.2005.03.010. PubMed PMID: 15890530.

20. Lander GC, Stagg SM, Voss NR, Cheng A, Fellmann D, Pulokas J, Yoshioka C, Irving C, Mulder A, Lau PW, Lyumkis D, Potter CS, Carragher B. Appion: an integrated, database-driven pipeline to facilitate EM image processing. J Struct Biol. 2009;166(1):95–102. doi: 10.1016/j.jsb.2009.01.002. PubMed PMID: 19263523; PMCID: PMC2775544.

21. Zheng SQ, Palovcak E, Armache JP, Verba KA, Cheng Y, Agard DA. MotionCor2: anisotropic correction of beam-induced motion for improved cryo-electron microscopy. Nat Methods. 2017;14(4):331–2. Epub 20170227. doi: 10.1038/nmeth.4193. PubMed PMID: 28250466; PMCID: PMC5494038.

22. Punjani A, Rubinstein JL, Fleet DJ, Brubaker MA. cryoSPARC: algorithms for rapid unsupervised cryo-EM structure determination. Nat Methods. 2017;14(3):290–6. Epub 20170206. doi: 10.1038/nmeth.4169. PubMed PMID: 28165473.

23. Punjani A, Zhang H, Fleet DJ. Non-uniform refinement: adaptive regularization improves single-particle cryo-EM reconstruction. Nat Methods. 2020;17(12):1214–21. Epub 20201130. doi: 10.1038/s41592-020-00990-8. PubMed PMID: 33257830.

24. Liebschner D, Afonine PV, Baker ML, Bunkoczi G, Chen VB, Croll TI, Hintze B, Hung LW, Jain S, McCoy AJ, Moriarty NW, Oeffner RD, Poon BK, Prisant MG, Read RJ, Richardson JS, Richardson DC, Sammito MD, Sobolev OV, Stockwell DH, Terwilliger TC, Urzhumtsev AG, Videau LL, Williams CJ, Adams PD. Macromolecular structure determination using X-rays, neutrons and electrons: recent developments in Phenix. Acta Crystallogr D Struct Biol. 2019;75(Pt 10):861–77. Epub 2019/10/08. doi: 10.1107/S2059798319011471. PubMed PMID: 31588918; PMCID: PMC6778852.

25. Emsley P, Lohkamp B, Scott WG, Cowtan K. Features and development of Coot. Acta Crystallogr D Biol Crystallogr. 2010;66(Pt 4):486–501. Epub 2010/04/13. doi: S0907444910007493 [pii] 10.1107/S0907444910007493. PubMed PMID: 20383002; PMCID: 2852313.

26. The PyMOL Molecular Graphics System. 2.5.2 ed: Schrödinger, LLC. Meng EC, Goddard TD, Pettersen EF, Couch GS, Pearson ZJ, Morris JH, Ferrin TE. UCSF ChimeraX: Tools for structure building and analysis. Protein Sci. 2023;32(11):e4792. doi: 10.1002/pro.4792. PubMed PMID: 37774136; PMCID: PMC10588335.

27. Case DA, Aktulga HM, Belfon K, Cerutti DS, Cisneros GA, Cruzeiro VWD, Forouzesh N, Giese TJ, Götz AW, Gohlke H, Izadi S, Kasavajhala K, Kaymak MC, King E, Kurtzman T, Lee TS, Li P, Liu J, Luchko T, Luo R, Manathunga M, Machado MR, Nguyen HM, O’Hearn KA, Onufriev AV, Pan F, Pantano S, Qi R, Rahnamoun A, Risheh A, Schott-Verdugo S, Shajan A, Swails J, Wang J, Wei H, Wu X, Wu Y, Zhang S, Zhao S, Zhu Q, Cheatham TE 3rd, Roe DR, Roitberg A, Simmerling C, York DM, Nagan MC, Merz KM Jr. AmberTools. J Chem Inf Model. 2023 Oct 23;63(20):6183–6191. doi: 10.1021/acs.jcim.3c01153. Epub 2023 Oct 8. PMID: 37805934; PMCID: PMC10598796.

28. Bergonzo C, Cheatham TE 3rd. Improved Force Field Parameters Lead to a Better Description of RNA Structure. J Chem Theory Comput. 2015 Sep 8;11(9):3969–72. doi: 10.1021/acs.jctc.5b00444. Epub 2015 Aug 18. PMID: 26575892.

29. Zgarbová M, Otyepka M, Sponer J, Mládek A, Banáš P, Cheatham TE 3rd, Jurečka P. Refinement of the Cornell et al. Nucleic Acids Force Field Based on Reference Quantum Chemical Calculations of Glycosidic Torsion Profiles. J Chem Theory Comput. 2011 Sep 13;7(9):2886–2902. doi: 10.1021/ct200162x. Epub 2011 Aug 2. PMID: 21921995; PMCID: PMC3171997.

30. Maier JA, Martinez C, Kasavajhala K, Wickstrom L, Hauser KE, Simmerling C. ff14SB: Improving the Accuracy of Protein Side Chain and Backbone Parameters from ff99SB. J Chem Theory Comput. 2015 Aug 11;11(8):3696–713. doi: 10.1021/acs.jctc.5b00255. Epub 2015 Jul 23. PMID: 26574453; PMCID: PMC4821407.

31. Li P, Song LF, Merz KM Jr. Parameterization of highly charged metal ions using the 12-6-4 LJ-type nonbonded model in explicit water. J Phys Chem B. 2015 Jan 22;119(3):883–95. doi: 10.1021/jp505875v. Epub 2014 Sep 12. PMID: 25145273; PMCID: PMC4306492.

32. Darden T, Perera L, Li L, Pedersen L. New tricks for modelers from the crystallography toolkit: the particle mesh Ewald algorithm and its use in nucleic acid simulations. Structure. 1999 Mar 15;7(3):R55–60. doi: 10.1016/s0969-2126(99)80033-1. Epub: 10368306.

33. Berendsen HJC, Postma JPM, van Gunsteren WF, DiNola A, Haak JR. Molecular dynamics with coupling to an external bath. J Chem Phys 1984. Oct 14;81(8):3684–90 doi: 10.1063/1.448118

34. Hopkins CW, Le Grand S, Walker RC, Roitberg AE. Long-Time-Step Molecular Dynamics through Hydrogen Mass Repartitioning. J Chem Theory Comput. 2015 Apr 14;11(4):1864–74. doi: 10.1021/ct5010406. Epub 2015 Mar 30. PMID: 26574392.

35. Roe DR, Cheatham TE 3rd. PTRAJ and CPPTRAJ: Software for Processing and Analysis of Molecular Dynamics Trajectory Data. J Chem Theory Comput. 2013 Jul 9;9(7):3084–95. doi: 10.1021/ct400341p. Epub 2013 Jun 25. PMID: 26583988.

36. Humphrey W, Dalke A, Schulten K. VMD: visual molecular dynamics. J Mol Graph. 1996 Feb;14(1):33–8, 27-8. doi: 10.1016/0263-7855(96)00018-5. Epub: 8744570.

37. Barnwal RP, Yang F, Varani G. Applications of NMR to structure determination of RNAs large and small. Arch Biochem Biophys. 2017 Aug 15;628:42–56. doi: 10.1016/j.abb.2017.06.003. Epub 2017 Jun 16. PMID: 28600200; PMCID: PMC5555312.

38. Jones AN, Tikhaia E, Mourão A, Sattler M. Structural effects of m6A modification of the Xist A-repeat AUCG tetraloop and its recognition by YTHDC1. Nucleic Acids Res. 2022 Feb 28;50(4):2350–2362. doi: 10.1093/nar/gkac080. Epub: 35166835; PMCID: PMC8887474.

39. Skinner SP, Fogh RH, Boucher W, Ragan TJ, Mureddu LG, Vuister GW. CcpNmr AnalysisAssign: a flexible platform for integrated NMR analysis. J Biomol NMR. 2016 Oct;66(2):111–124. doi: 10.1007/s10858-016-0060-y. Epub 2016 Sep 23. Erratum in: J Biomol NMR. 2017 Apr;67(4):321. doi: 10.1007/s10858-017-0102-0. Epub: 27663422; PMCID: PMC5095159.

