## Supplementary Information for "Structural and biophysical insights into the unique RNase YicC"

**Table S1.**
**Oligonucleotides used in this study**

| Name | Sequence |
|---|---|
| 2051 | 5'-GGCAGAAGAAUGCUGUAAAACAGAGA-3' |
| 2176 | 5'-GGAAGAAGAAUGCUGUAAAACAGAGA -3' |
| 2182 | 5'-pGGCAGAAGAAUGCUGUAAAACAGAGA-3' |
| 2292 | 5'-AGAGACAAAAUGUCGUAAGAAGACGG -3' |
| 2305 | 5'-AGAGACAAAAUGAGCUAAGAAGACGG -3' |
| 2051A | 5'-GGCAGAAGAAUGCU-3'[a] |
| 2051B | 5'-AGAAUGCUGUAAAACAGAGA-3' |
| 1935 | 5'-GGCGAUGCUGUAACGCGGCGAUAAGU-3' |
| 1955 | 5'-ACUUAUCGCCGCGUUACAGCAUCGCC-3' |

[a]Note: For IVT experiments an extra G was added to the 5' end of 2051A

**Table S2. Cryo-EM data collection, refinement and validation statistics**

| | YicC E281A-2182 | YicC E281A-2176 with $Mg^{2+}$ | YicC E281A-2051 |
|---|---|---|---|
| **Data collection and processing** | | | |
| Magnification | 105,000 | 81000 | 81,000 |
| Voltage (kV) | 300 | 300 | 300 |
| Electron exposure (e–/Å$^2$) | 54.03 | 55.95 | 51.50 |
| Defocus range (μm) | 0.16 to -2.68 | 0.6 – 1.8 | -0.8 to -2.0 |
| Pixel size (Å) | 0.826 | 0.8560 | 0.856 |
| Symmetry imposed | C1 | C1 | C1 |
| Initial particle images (no.) | 1,470,323 | 3,552,024 | 1,704,486 |
| Final particle images (no.) | 180,679 | 166,251 | 270,956 |
| Map resolution (Å) | 3.02 | 3.04 | 3.73 |
| FSC threshold | 0.143 | 0.5 | 0.143 |
| Map resolution range (Å) | 2.6 -11.9 | 2.5 – 6.28 | 3.11-6.25 |
| **Refinement** | | | |
| Initial model used (PDB code) | 8VES | 2182 structure | 8VES |
| Map sharpening *B* factor (Å$^2$) | -126.2 | -108.3 | -189.0 |
| Model composition | | | |
| Non-hydrogen atoms | 14870 | 15037 | 14282 |
| Protein residues | 1713 | 1713 | 1714 |
| Nucleotides | 31 | 36 | 19 |
| *B* factors (Å$^2$) | | | |
| Protein | 48.62 | 78.94 | 123.92 |
| Nucleotides | 32.15 | 47.68 | 99.77 |
| R.m.s. deviations | | | |
| Bond lengths (Å) | 0.008 | 0.011 | 0.008 |
| Bond angles (°) | 0.732 | 0.871 | 0.911 |
| Validation | | | |
| MolProbity score | 1.71 | 1.75 | 1.61 |
| Clashscore | 16.02 | 17.99 | 9.31 |
| Poor rotamers (%) | 0 | 0 | 0 |
| Ramachandran plot | | | |
| Favored (%) | 98.29 | 98.11 | 97.41 |
| Allowed (%) | 1.65 | 1.83 | 2.59 |
| Disallowed (%) | 0.06 | 0.06 | 0 |

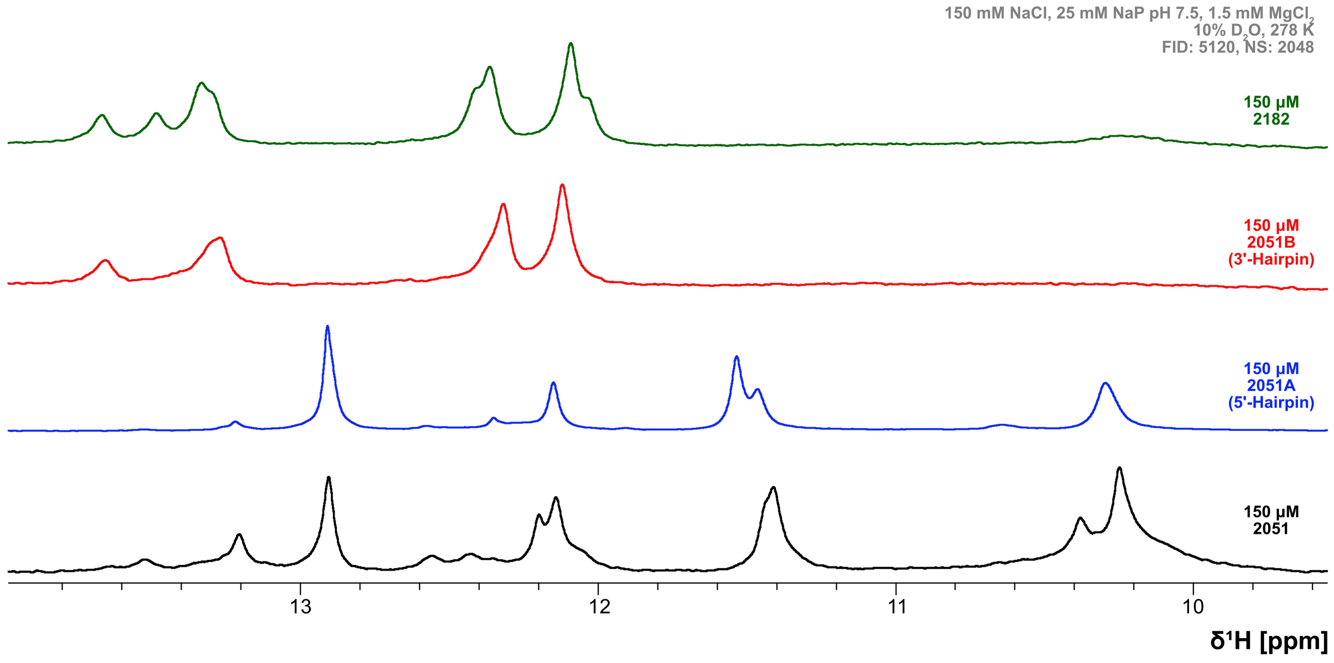


Figure S1. 1D 1H NMR spectra of RNA oligos in the presence of magnesium. All measurements were performed with identical buffer conditions at 278K.

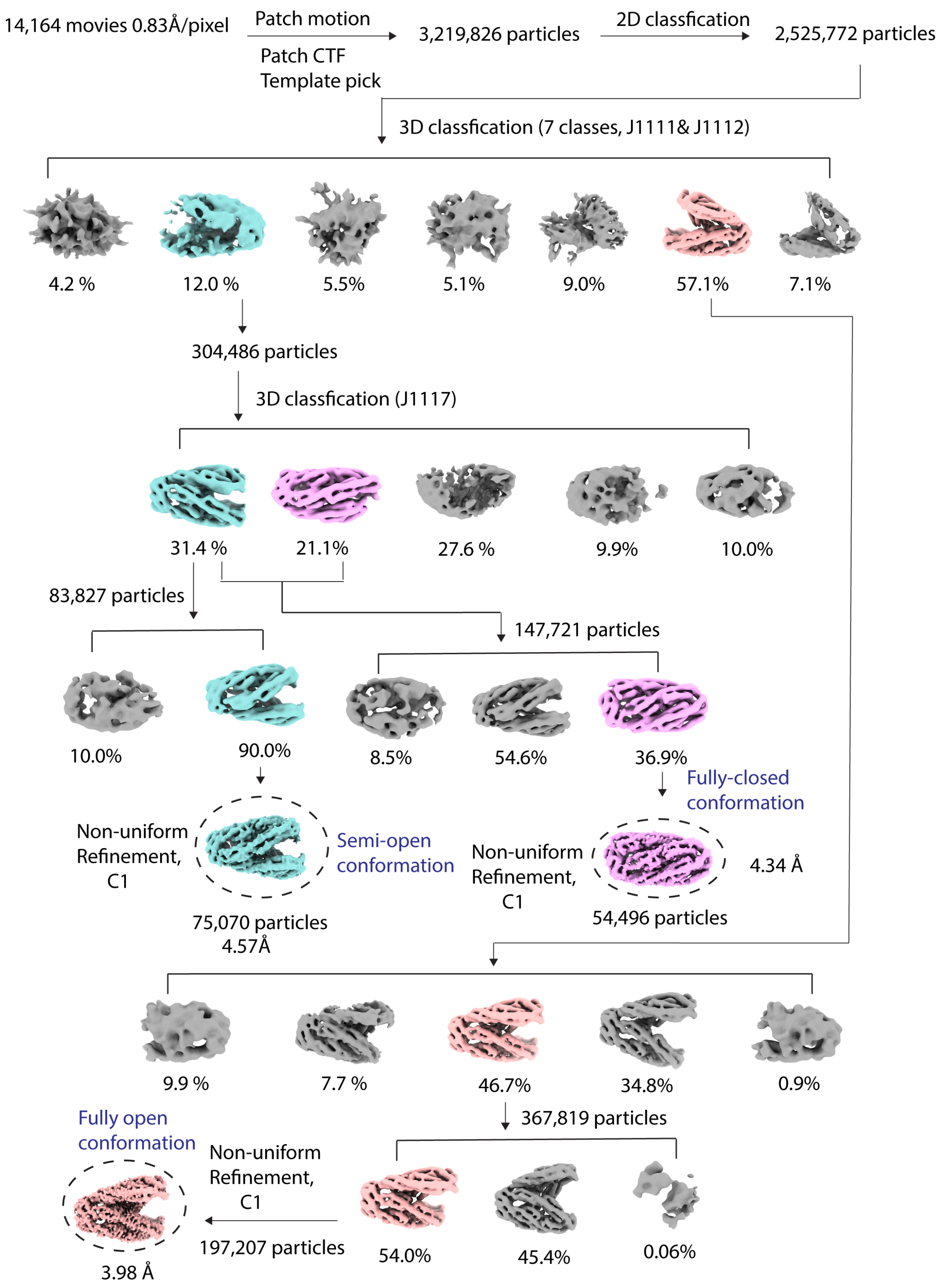


Figure S2. Cryo-EM work-flow for low-resolution WT:2176 complexes.

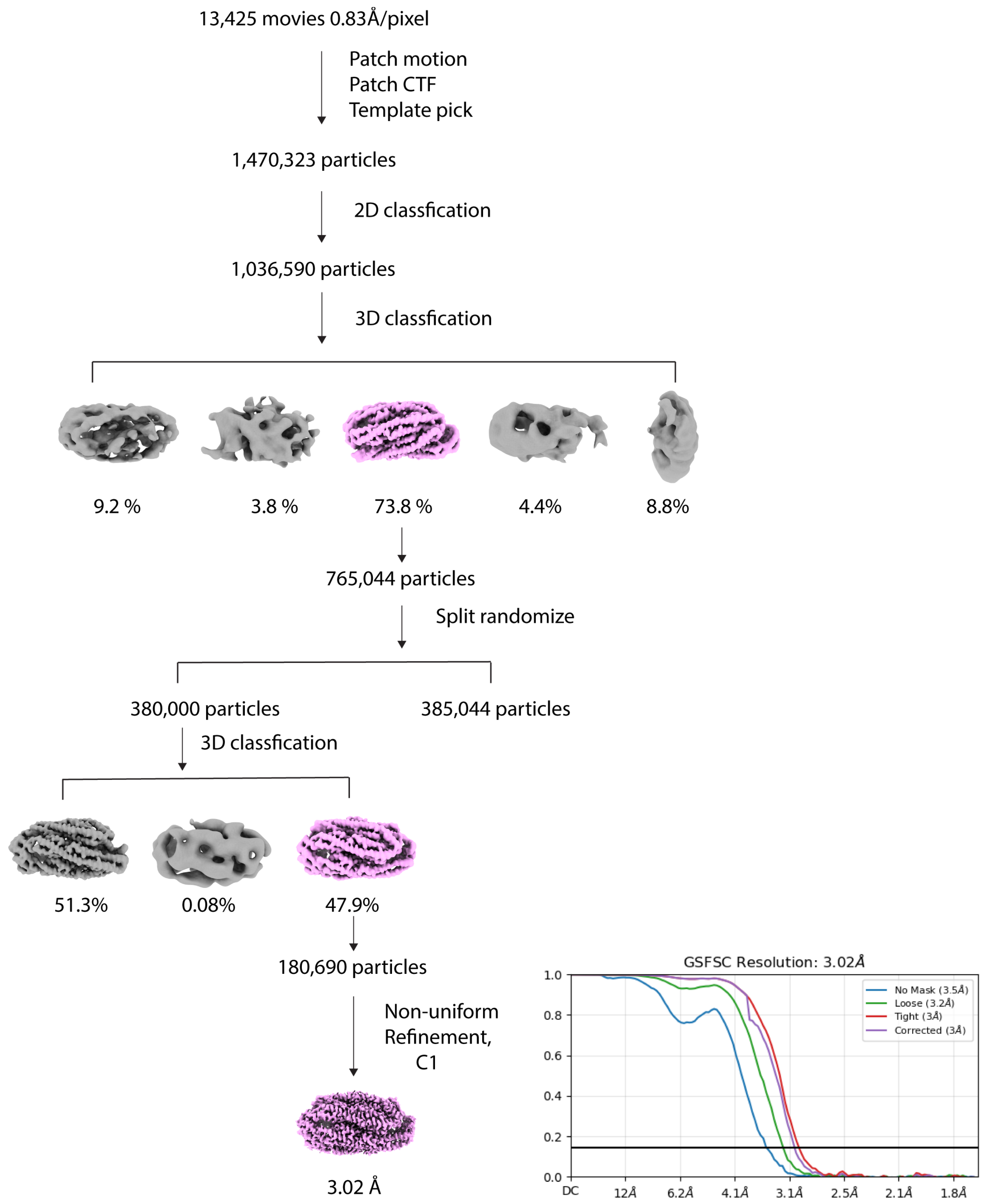


Figure S3. Cryo-EM work-flow for E281A:2182 complex.

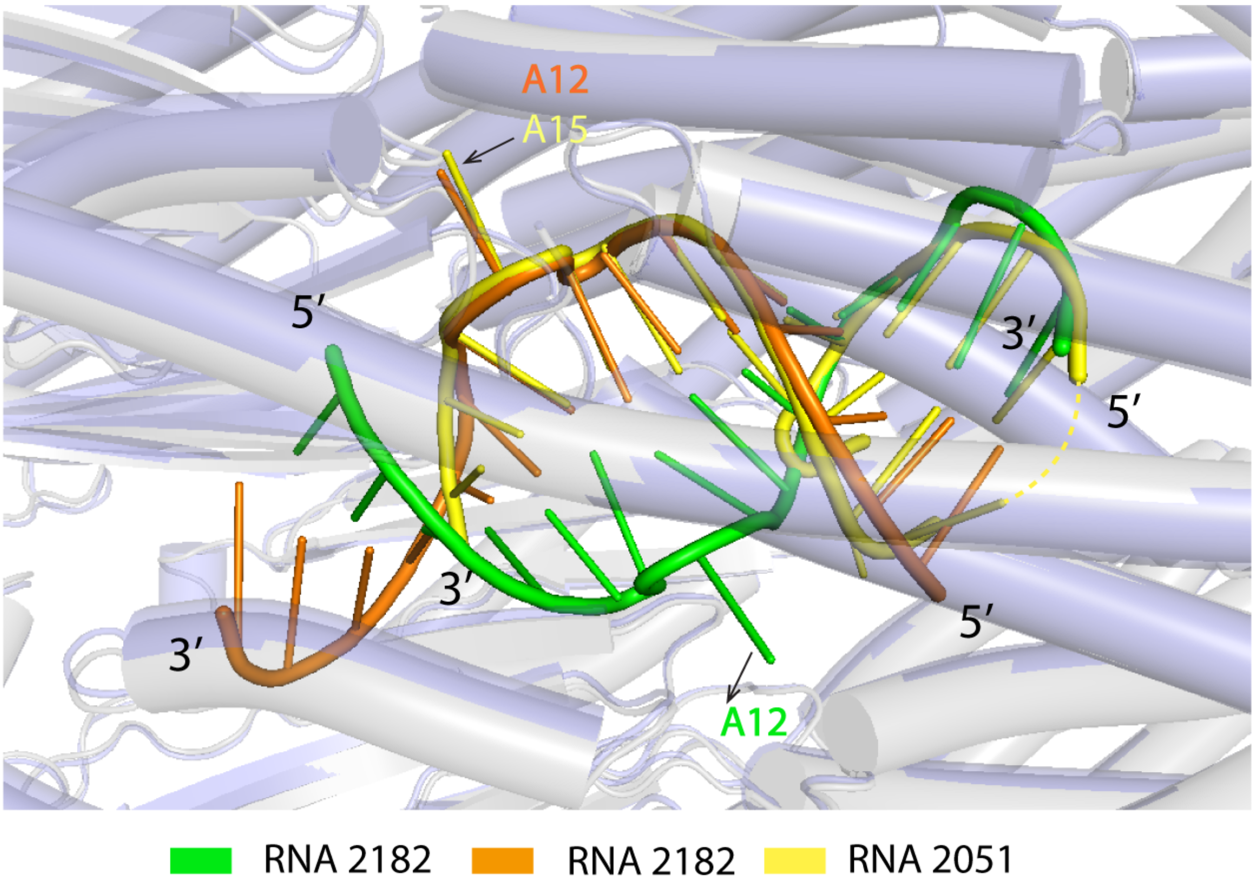


Figure S4. Overlay of WT:2051 structure and E281A:2182 structure. The 2051 RNA (from 8VES) is shown as a cartoon in yellow, and the dimer of 2182 is shown in green and orange. There is not density for the loop of the 2051 hairpin, so it is represented as a dotted line. The adenine that gets extruded from each strand is indicated, highlighting the identical overlap on the 2051 and 2182 top strand.

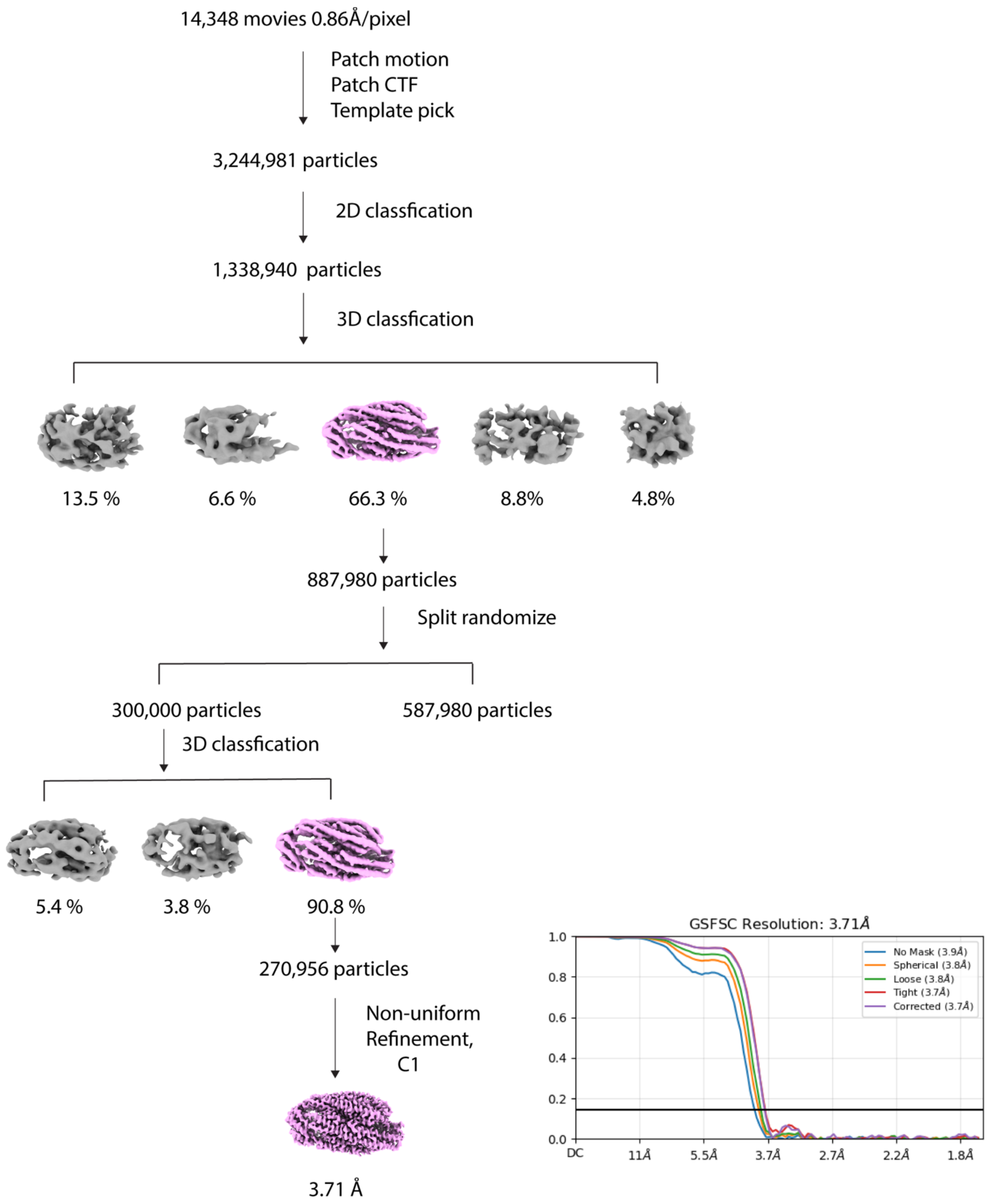


Figure S5. Cryo-EM work-flow for E281A:2051 complexes.

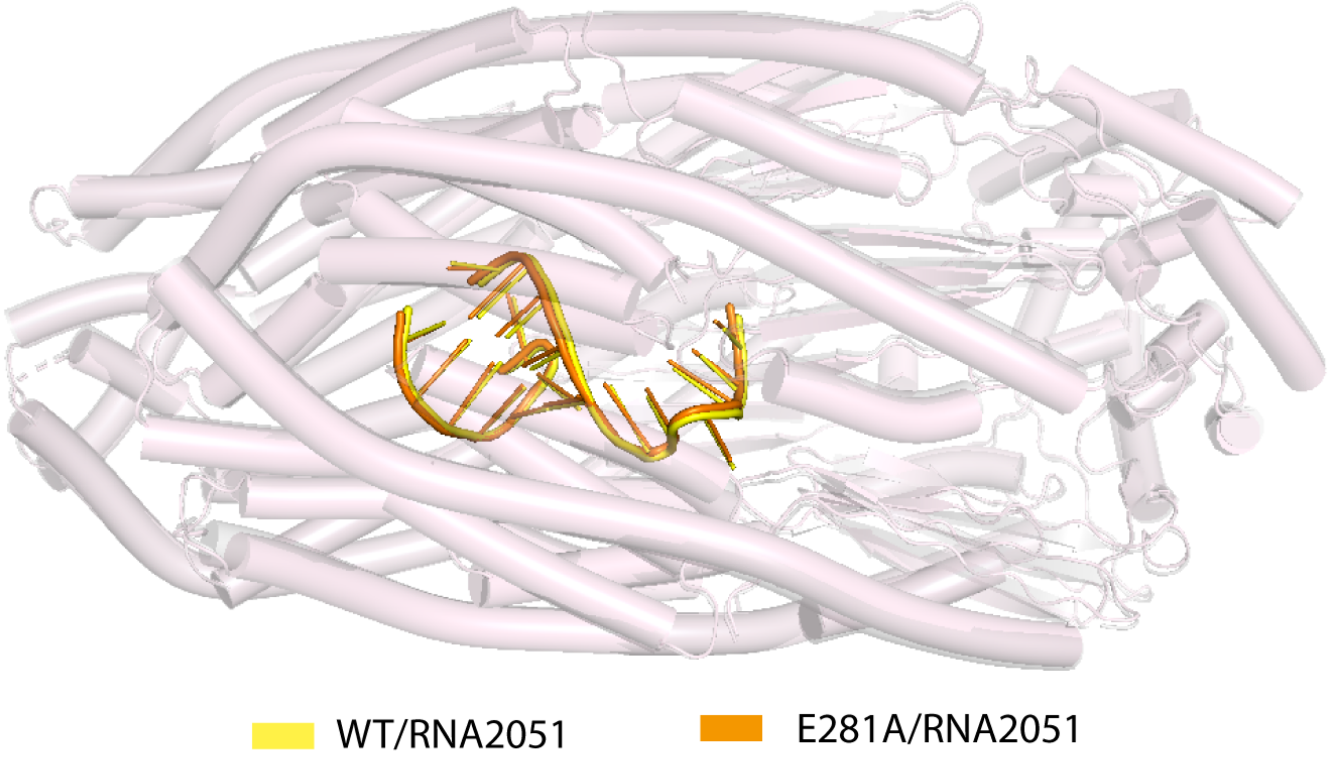


Figure S6. Overlay of wild-type and E281A mutant YicC structures bound to 2051. The protein is shown as a cartoon and the RNA is colored as shown.

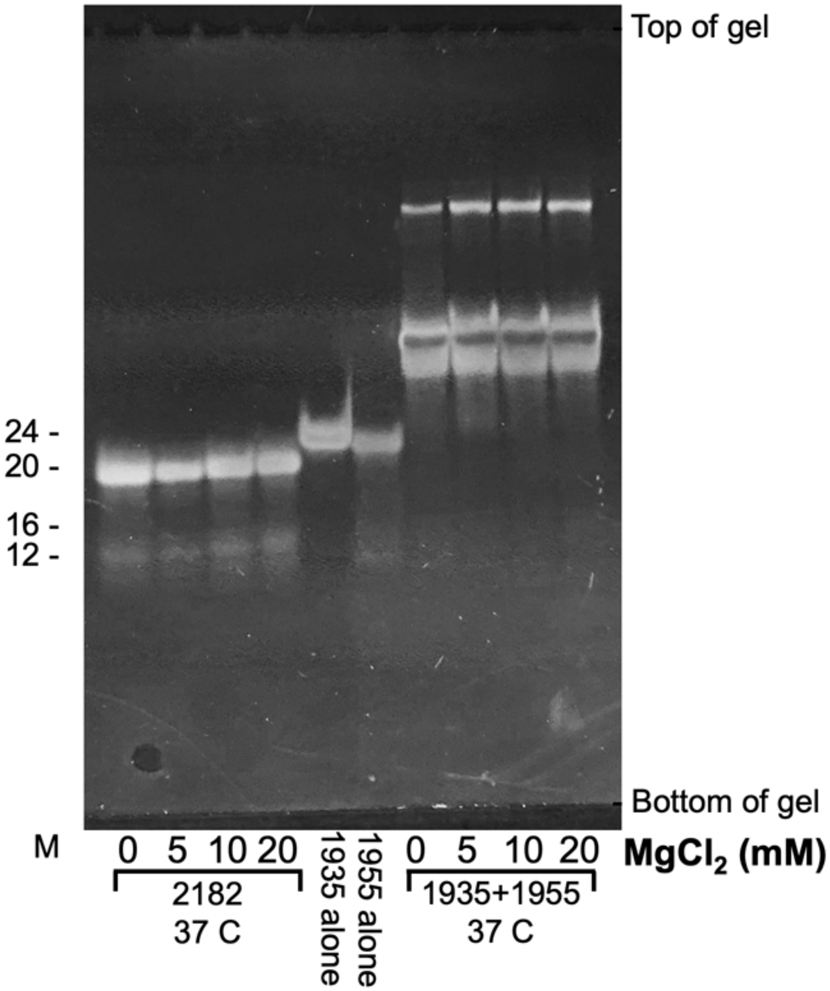


**Figure S7. Dimerization of RNA in solution**. Indicated RNA oligonucleotides were heated to 37°C in different concentrations of magnesium and assayed on a non-denaturing 15% polyacrylamide gel for dimer formation. Migration on this gel of single-stranded oligos of the sizes shown is indicated at the left.

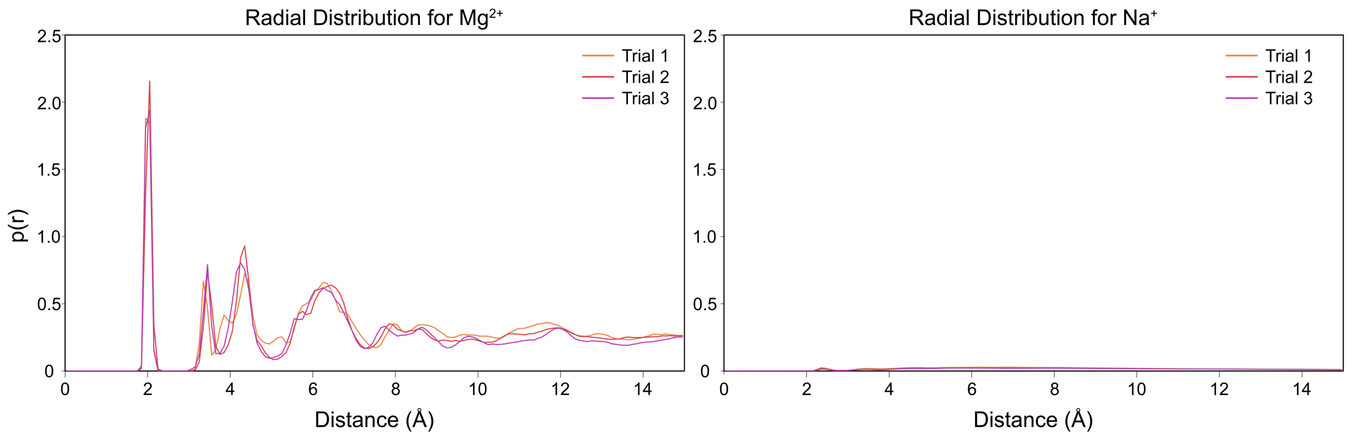


Figure S8. Radial distribution function plots for the 8VES system (WT YicC system bound to 2051 RNA oligo) comparing $Mg^{2+}$ and $Na^{2+}$ density around heavy atoms within the proposed cleavage site.

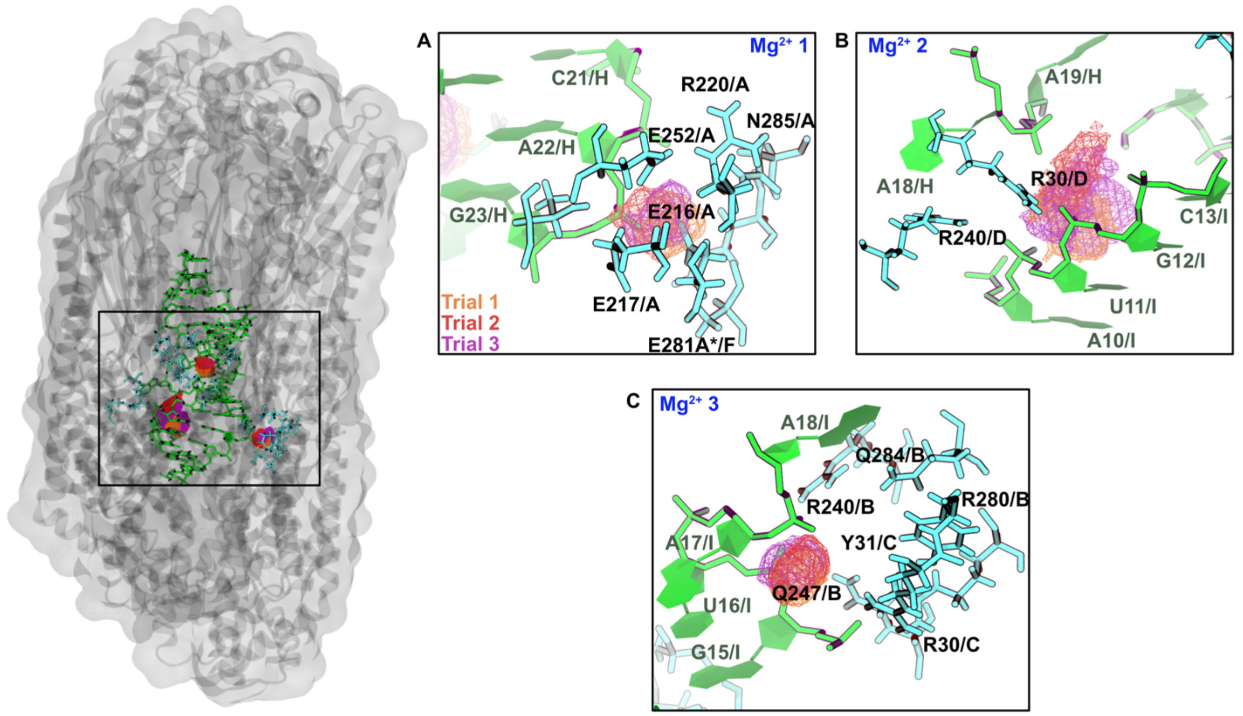


Figure S9. Molecular dynamics simulations of E281A bound to 2182. Ion density mapping was performed on each $Mg^{2+}$ ion over the course of the simulation. Three possible sites of magnesium density are highlighted. (A) First localization occupies space around a glutamate pair (E216/A, E217/A). (B) Second $Mg^{2+}$ ion competes with arginine residues (R230/D, R240/D) to coordinate with the RNA backbone. (C) Third $Mg^{2+}$ localization shows preference to RNA pocket, sequestered away from protein residues.

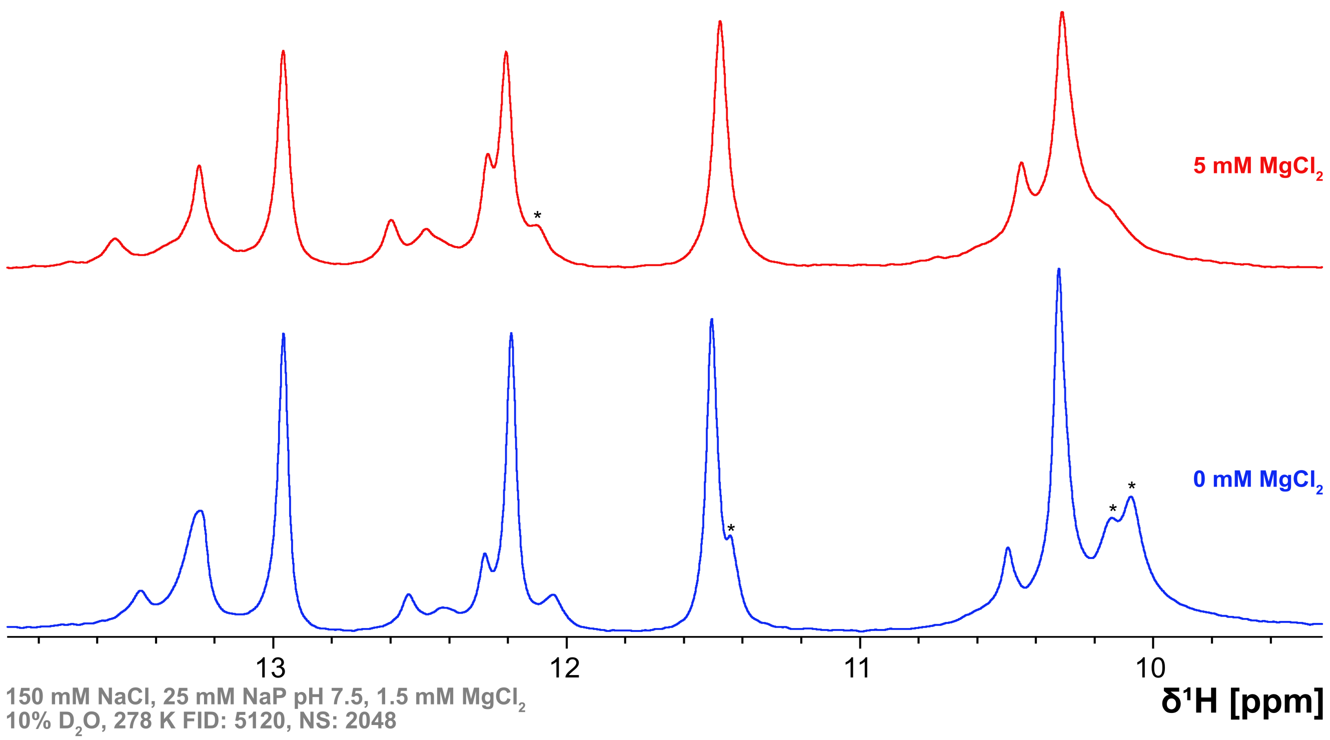


Figure S10. 1D-[1]H NMR spectra of FL-2051 RNA in either the presence (red) or absence (blue) of $MgCl_2$. Spectral differences are highlighted with asterisks, with slight chemical shift differences observed upfield around 10.1 ppm.

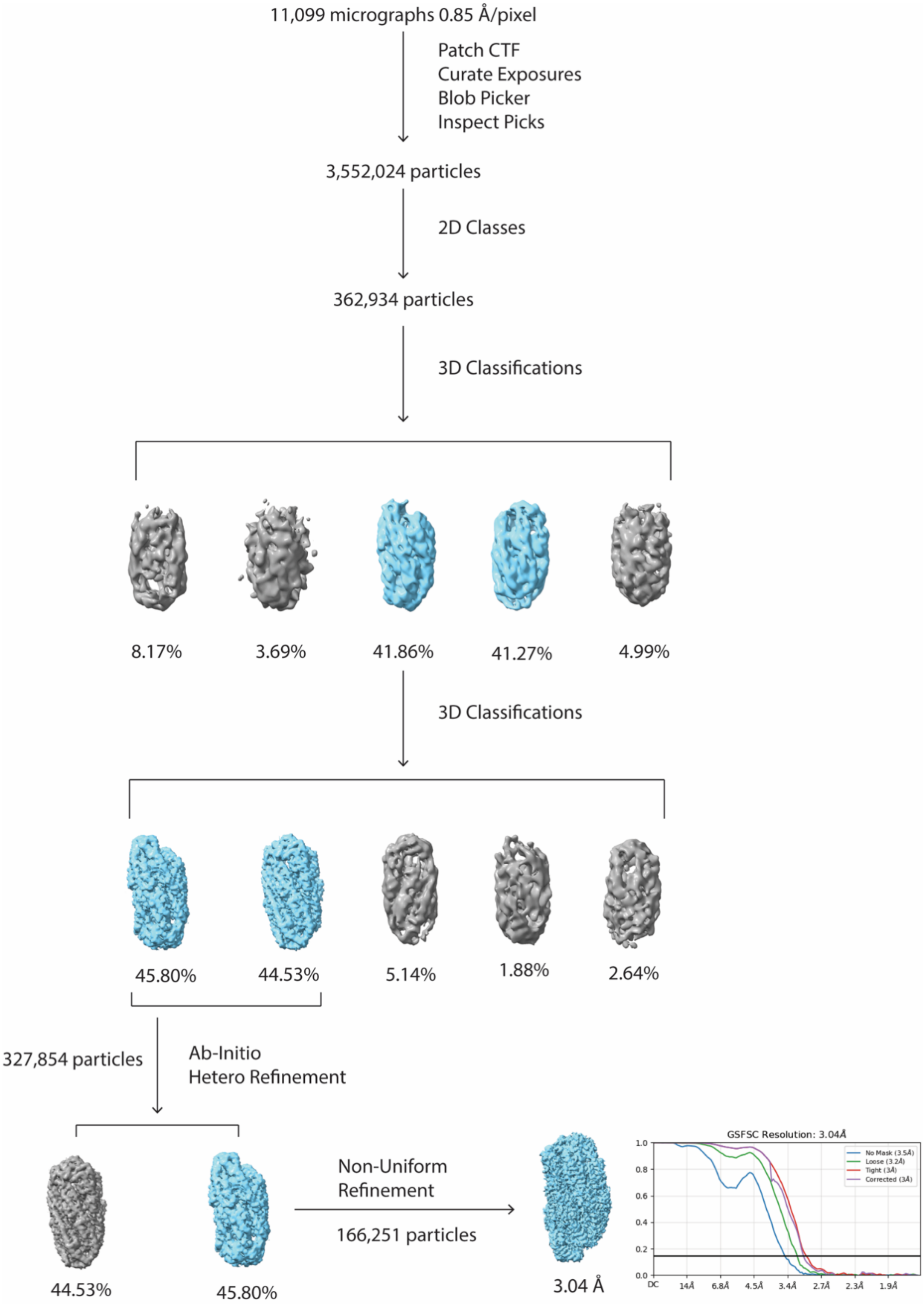


Figure S11. Cryo-EM work-flow for E281A:2176 complexes with $Mg^{2+}$.

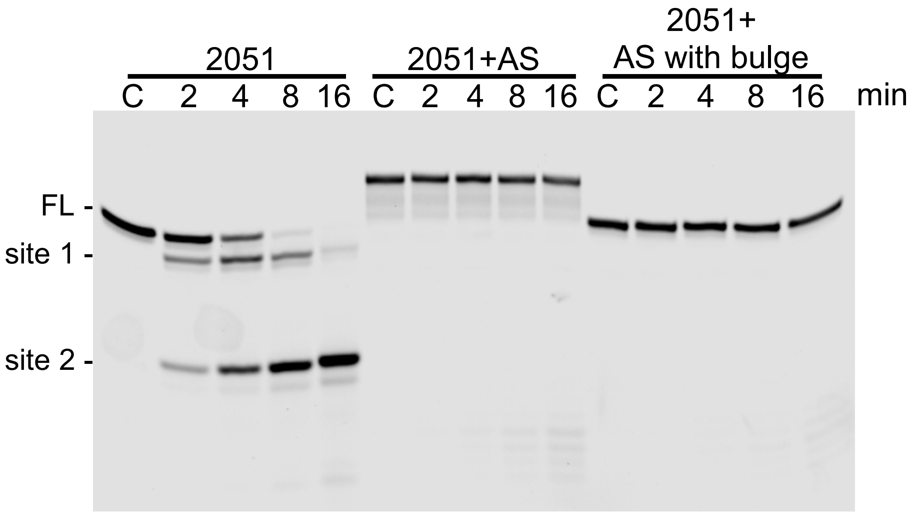


5'-GGCAGAAGAAUGCUGUAAAACAGAGA-3'
3'-CCGUCUUCUUACGACAUUUUGUCUCU-5'
2051:2292

$\quad\quad\quad\quad\quad\quad\quad$ G C U
5'-GGCAGAAGAAU GUAAAACAGAGA-3'
3'-CCGUCUUCUUA UCAUUUUGUCUCU-5'
$\quad\quad\quad\quad\quad\quad\quad$ G C
2051:2305

Figure S12. Denaturing gel electrophoresis (20% polyacrylamide) showing YicC cleavage products of 3'-end labeled oligo 2051, in the absence and presence of antisense RNA. Ratio of antisense:sense RNA was 1.25:1. Migration of full-length (FL) oligo and products of cleavage at sites 1 and 2 are indicated at left. Time in min indicated above each lane. Control lanes (C) contained oligo with no YicC added. Below are the schematics of the duplexes.

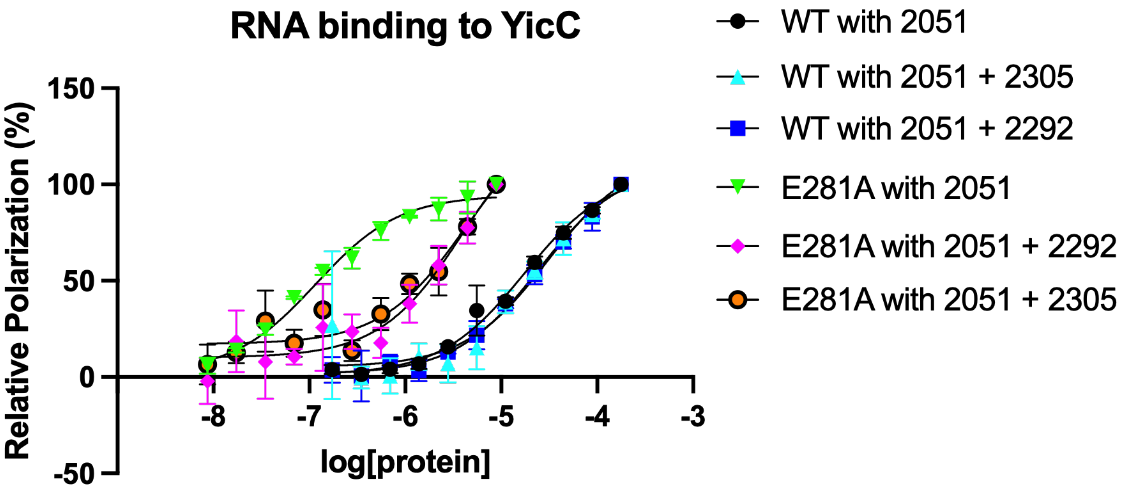


Figure S13. Binding of RNA duplexes to YicC protein. Fluorescently labeled RNAs (3' fluorescein) were measured for polarization with varying protein concentrations. n=3 and error bars indicate SD.

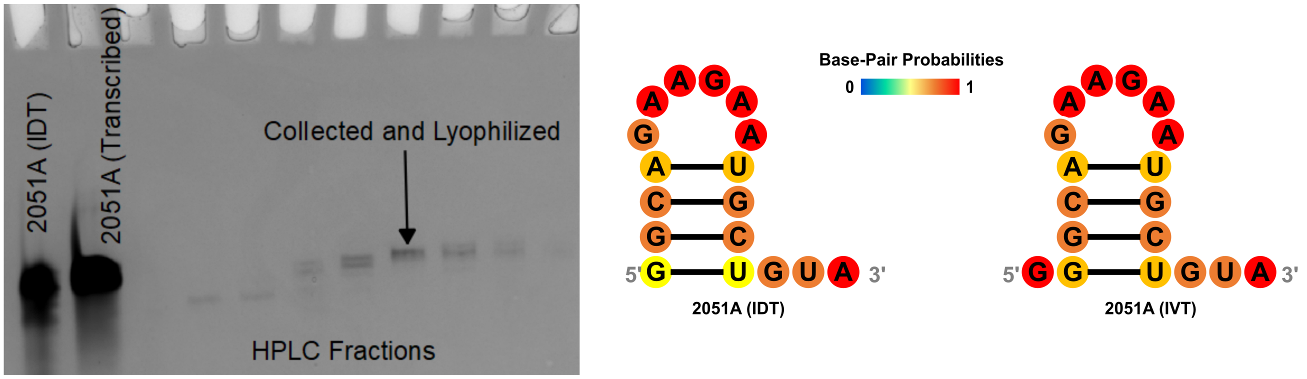


Figure S14. 20% denaturing polyacrylamide gel of transcribed 2051A. Gel shows HPLC purification of IVT transcribed 2051A compared with 2051A which was purchased from IDT. The designed transcript expectedly is one nucleotide larger to improve transcription efficiency with T7 polymerase. No structural changes are predicted with ViennaRNA upon incorporation of this additional G nucleotide at the 5’ site.
